# Stringent proteogenomic discovery of novel small proteins in *Mycobacterium tuberculosis* clinical reference strains

**DOI:** 10.64898/2026.01.27.701740

**Authors:** Benjamin Heiniger, Christian Schori, Mohammad Arefian, Amir Banaei-Esfahani, Martin Schuler, Sonia Borrell, Chloé Loiseau, Daniela Brites, Iñaki Comas, Ruedi Aebersold, Sebastien Gagneux, Ben C. Collins, Christian H. Ahrens

## Abstract

Even though our meta-analysis ranks *Mycobacterium tuberculosis* genomes among the bacterial pathogens that are most straightforward to assemble, most available assemblies relied on short-read sequencing and contain genomic blind spots that miss functionally important genes. Complete genomes are essential for functional genomics, particularly for identifying small ORF-encoded proteins (SEPs; ≤100 amino acids), which can play critical biological roles yet are frequently missed by standard annotations. Here, we generated complete long-read assemblies for six clinical reference strains representing lineage 1 and the more pathogenic lineage 2, followed by comparative genomic and proteogenomic analyses. We additionally provide software to predict comprehensive sets of mycobacteria-specific proline-glutamic acid (PE) and PPE family genes, including lineage-specific variants.

Using parallel accumulation–serial fragmentation mass spectrometry, we detected approximately two-thirds of each strain’s annotated proteome from unfractionated cell extracts. Extending our proteogenomic framework across related strains, and adding rigorous control of proteogenomic discovery rates using entrapment strategies, we revealed 12–24 previously unannotated proteins per strain, predominantly SEPs, 56–60 alternative translation start sites, and 9–17 expressed pseudogenes. Newly identified proteins included conserved and lineage-specific SEPs, an antitoxin, candidate antimicrobial peptides and novel proteins under purifying selection. Overall, applying this improved proteogenomics method to phylogenomically selected clinical reference strains provides a valuable approach for discovering candidate diagnostics or therapeutics, as illustrated here for a WHO-listed critical bacterial pathogen.

## Introduction

Strains of the human-adapted *Mycobacterium tuberculosis* complex (MTBC) are among the deadliest bacterial pathogens, claiming 1.23 million lives in 2024^5^. Thousands of MTBC strains have been sequenced, their phylogenies reconstructed and ten human-adapted MTBC lineages (L) identified^1^. They encompass strains of the ancestral L1 and L5-10, and of the modern, more virulent lineages 2, 3 and 4, that differ in the presence of the TbD1 genomic region^2^. L1-L4 are globally most frequent, while L5-L10 are rarer and restricted to different regions of Africa^1^. With increasing multidrug resistance posing a major threat^3^, the wealth of genomic data is expected to offer avenues to identify novel therapeutic solutions^4,5^. Twenty clinical reference strains were previously selected to capture the diversity within and between seven of the ten major human-adapted MTBC lineages^6^, i.e., L1-L7. They are a valuable resource to move beyond studying the MTBC based on laboratory strains (mainly from L4), particularly for multi-omics^7^ and functional studies. Due to their clinical relevance, strains from L1 and L2 have gained particular interest. Strains from L1 cause the highest TB burden in terms of absolute number of cases occurring globally^1^, but exhibit a lower virulence in infection models, while L2 strains are linked with higher virulence, faster disease progression and positively associated with antibiotic resistance^8^. As L1 and L2 do not comprise other species, we use the term Mtb from here on.

Most bacterial genomes have been assembled from short read data which cannot resolve longer repetitive regions, mobile genetic elements and structural rearrangements^9,10^. Such assemblies can miss important genes with roles in antibiotic resistance (non-ribosomal peptide synthetases), virulence and immune evasion (adhesins) and killing competitors (type VI secretion system effectors) and even essential genes as shown for *Pseudomonas aeruginosa* MPAO1, a model of another highly relevant pathogen^11^. The *M. tuberculosis* PE and PPE gene families, named after conserved proline-glutamic acid (PE) and proline-proline-glutamic acid (PPE) residues in their N-terminus^12,13^, are exclusive to the genus Mycobacteria^14^ and contain many repeat sequences that are difficult to resolve with short read data. In contrast to the reductive genome evolution observed in Mtb^15^, these gene families have greatly expanded, attesting to their importance for the success of Mtb strains in their ecological niche^16^. Despite their relevance for virulence, host-pathogen interaction and as potential targets for the development of novel drugs, diagnostics and vaccines^17^, these genes have thus long represented genomic blind spots^18^. The advantages of long read sequencing for complete de novo genome assembly of repeat-rich bacterial genomes have been illustrated^19^ and recently also applied to Mtb^20^. Complete genomes are the ideal basis for functional and comparative genomics^19,21^.

Complete genomes can be interrogated with proteomics or ribosome profiling to uncover evidence for so far unannotated genes, many of which encode small ORF-encoded proteins (SEPs)^22–24^. SEPs can carry out important functions, e.g. in cell-to-cell communication^25^, membrane complex assembly, regulation of protein stability and/or activity^26^, have roles in antibiotic resistance^27^ and as antimicrobial peptides (AMPs) they are able to shape microbial community composition^28^. Yet, the small proteome has been notoriously understudied due to missed coding sequences (CDS), the small number of tryptic peptides produced and as modifications are required for each step of the analytical workflow to increase the detection rate by mass spectrometry (MS)^29^. Ribosome profiling (Ribo-seq) captures actively translated mRNAs^24^ and is highly sensitive, but needs to be adapted to each bacterial model system. Advances in MS workflows such as top-down proteomics^30^ and more sensitive bottom-up workflows such as parallel accumulation-serial fragmentation (PASEF)^31^, which can acquire spectra at a higher rate, are alternative options to more comprehensively cover the small proteome in a given condition.

Mining public meta-data, we here assess the suitability of Mtb strains for genomic and proteogenomic studies. Complete genomes were assembled for three clinical reference strains from L1 and L2 each, allowing to report complete inventories of their PE and PPE gene families, including software to create them for any Mtb full genome, and to perform comparative genomics. Adapting our proteogenomics approach for the analysis of phylogenomically selected strains of interest, we identified three types of novelties found in all six strains, in a specific lineage or in a subset of the strains: novel CDS (mostly SEPs below 100 aa), longer or shorter proteins than annotated by RefSeq and expressed pseudogenes. The novel SEPs included several interesting examples: an antitoxin, potential antimicrobial peptides (AMPs) and several novel ORFs under purifying selection. The false discovery proportion (FDP) estimate generated by entrapment^32^, and evaluated in partitioned canonical RefSeq compared to novel identifications, allowed us to more rigorously control the higher local false discovery rate (FDR) among novel proteins—an issue that is often inadequately addressed in proteogenomics and has even been termed the Achilles heel of proteogenomics^71^.

## Methods

### Meta-analysis of *M. tuberculosis* genomes and assembly complexity

The assembly level of all Mtb genomes in the National Center of Biotechnology Information’s (NCBI) RefSeq database (DB) (Dec 31, 2024), was parsed from the downloaded summary file (https://ftp.ncbi.nlm.nih.gov/genomes/refseq/bacteria/assembly_summary.txt). The number of contigs per assembly was extracted (column contig_count); the count of long read (PacBio or ONT), hybrid and short read only-based assemblies (Illumina, etc.) (**Supplementary Fig. S1**) was parsed from the sequencing technology field of the downloaded Genbank files with BioPython 1.76. The genome assembly complexity is a metric that classifies prokaryotic genomes based on their total number of repeats (≥ 500 bp, with a sequence identity ≥ 95%) and the length of the longest repeat into three classes^33^: easy to assemble class I genomes (up to 100 repeats, the longest repeat is the entire rRNA operon of up to 6.5-7kb), class II genomes (more than 100 repeats, longest repeat rRNA operon) which e.g. include various *Lactobacillales*, which often undergo reductive genome evolution and can harbor hundreds of transposons, and difficult to assemble class III genomes whose repeats can be much longer and extend even well over 100kb^19^. The genome assembly complexity was computed for the six Mtb strains and for all complete RefSeq genomes of selected bacterial pathogens, i.e., critical and top priority pathogens listed by the WHO^70^ and from^34^.

### Cell culture, DNA extraction, sequencing, assembly and annotation

Six Mtb isolates were selected from a set of 20 clinical reference strains^6^, three from L1 (N0072 (ITM-500942; CT2018-00083), N0153 (ITM-500943; CT2018-00084), N0157 (ITM-500941; CT2018-00082)) and three from L2 (N0052 (ITM-601826; CT2018-03241), N0145 (ITM-500944; CT2018-00085), N0155 (ITM-500946; CT2018-00088)). Bacteria were grown in modified 7H9 medium supplemented with 0.5% w/v pyruvate, 0.05% v/v tyloxapol, 0.2% w/v glucose, 0.5% bovine serum albumin (Fraction V, Roche) and 14.5 mM NaCl (while tween 80, oleic acid, glycerol and catalase were omitted). For global protein expression profiling experiments, bacteria were cultured in 1l flasks at 37°C containing 100 ml media and large glass beads to avoid clumping and were rotated on a roller. Genomic DNA (gDNA) for Illumina and Pacific Biosciences’ (PacBio) sequencing was extracted as described (**Supplementary Methods**). *De novo* genome assemblies of the six strains were created with long read sequencing data from PacBio^7^, essentially as described previously^35^; for more detail, see **Supplementary Methods**. Assemblies were annotated with NCBI’s Prokaryotic Genome Annotation Pipeline (PGAP) v.4.8^36^. For an overview of the genome assemblies and selected genome features, see **Supplementary Tables S1** and **S2**.

### Comparison of complete genomes and Illumina assemblies

De novo genome assemblies were also created for all six strains only using Illumina short reads (see **Supplementary Methods**). Contigs were mapped to the complete PacBio assemblies using minimap2 (version 2.17)^37^, missing genes were identified by comparison with PGAP annotated coding sequences (CDS) (partially missed genes: at least 5% of the CDS sequence were missed in the short-read assembly). For an overview of gene families enriched among missing/incomplete CDS, see **Supplementary Table 4**. Mapped Illumina contigs, gaps and missing genes were visualized together with sliding window means of the PacBio read coverage (**Fig. 2B**, **Supplementary Fig. S3**).

### Comparative genomics of clinical reference strains from L1, L2

An existing phylogenetic analysis^1^ was adapted to highlight the clinical reference strains from L1, L2, the laboratory model strains H37Rv, Erdman, CDC1551 from L4 and the clinical isolate HN878 (**Fig. 2A**). For the pangenome analysis, all non-pseudogene CDS without ribosomal frameshifts were analyzed using Panaroo 1.5.2^38^ without merging paralogs (parameters: --alignment pan, --clean-mode sensitive, --refind-mode off); the last two options allow to retain all genes even if only present in one strain (**Supplementary Table S5)**. Panaroo was chosen as it outperformed other tools due to its calculation of superior ortholog clusters and conservative pangenome size estimation^38^ and its better performance on Mtb genomes^21^. An UpSet plot (v0.9.0 of the Python package) was generated using the final presence-absence gene list (**Fig. 2C**), and for the PE/PPE families (**Supplementary Fig. S4**). The analysis was repeated with H37Rv included to characterize differences to this well-studied laboratory model strain^14^.

### Classification of PE and PPE family genes

For each strain, the set of PE and PPE family genes were predicted using three methods: The PGAP annotation was searched for a matching regular expression (regex), a reciprocal best BLAST hit (RBBH) was used to match all annotated PGAP proteins to a set of manually curated and annotated PE and PPE proteins from H37Rv^12^, and the descriptions of PFAM domains identified by an InterProScan analysis were searched with the same regex set. The first unambiguous assignment from this order was taken (for more detail, see **Supplementary Methods**). The PE and PPE family proteins were then further classified into one of five sublineages of the corresponding protein family by performing a phylogenetic analysis on their N-terminal sequences as described^13^ (**Table 1**). Finally, a motif-based subfamily prediction was used for those cases where the protein had no RBBH from which the annotated subfamily for the PE or PPE genes could be assigned. If the sequence contained a subfamily-specific minimum of motif matches, the protein was assigned to the PE-PGRS, PPE-PPW, PPE-SVP or PPE-MTRS subfamily; for more detail, see **Supplementary Methods, Supplementary Table S6**. As PE-PGRS subfamily proteins can only be secreted if an intact ppe38-71 locus is present, its configuration was analyzed for all six strains and compared to known configurations^12^ to assess its functionality (**Supplementary Fig. S5**). Since the corresponding genes were not annotated by PGAP for all six strains, orthologs of the H37Rv ppe38-71 genes were identified by tblastn. Finally, a Snakemake workflow to perform the PE/PPE classification for other Mtb isolates is made available as a resource to the community (**Data Availability**).

**Table 1.**
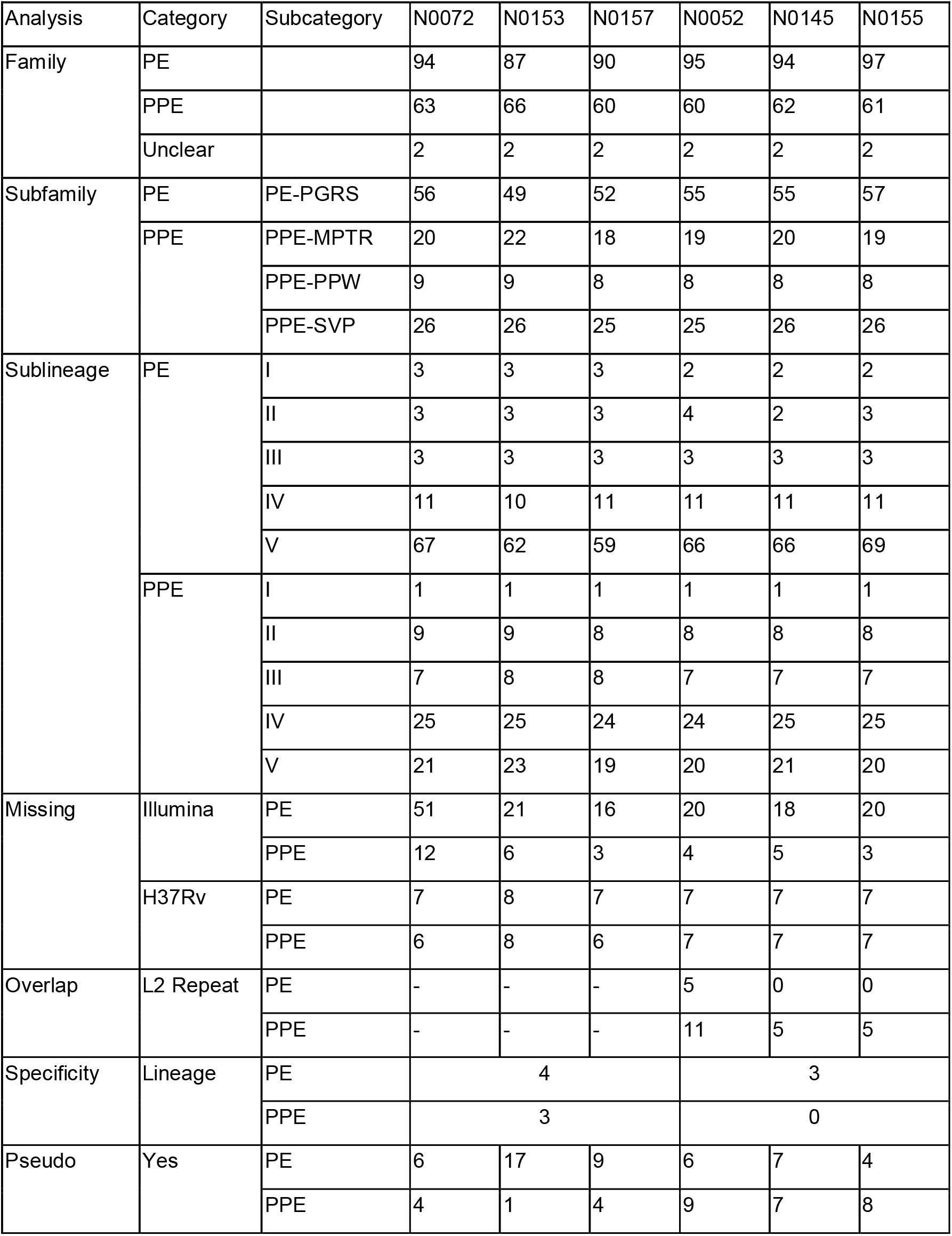
Catalog of PE/PPE gene family members and their sub-classification. For more detail, see Supplementary Table S6.

| Analysis | Category | Subcategory | N0072 | N0153 | N0157 | N0052 | N0145 | N0155 |
| --- | --- | --- | --- | --- | --- | --- | --- | --- |
| Family | PE |  | 94 | 87 | 90 | 95 | 94 | 97 |
|  | PPE |  | 63 | 66 | 60 | 60 | 62 | 61 |
|  | Unclear |  | 2 | 2 | 2 | 2 | 2 | 2 |
| Subfamily | PE | PE-PGRS | 56 | 49 | 52 | 55 | 55 | 57 |
|  | PPE | PPE-MPTR | 20 | 22 | 18 | 19 | 20 | 19 |
|  |  | PPE-PPW | 9 | 9 | 8 | 8 | 8 | 8 |
|  |  | PPE-SVP | 26 | 26 | 25 | 25 | 26 | 26 |
| Sublineage | PE | I | 3 | 3 | 3 | 2 | 2 | 2 |
|  |  | II | 3 | 3 | 3 | 4 | 2 | 3 |
|  |  | III | 3 | 3 | 3 | 3 | 3 | 3 |
|  |  | IV | 11 | 10 | 11 | 11 | 11 | 11 |
|  |  | V | 67 | 62 | 59 | 66 | 66 | 69 |
|  | PPE | I | 1 | 1 | 1 | 1 | 1 | 1 |
|  |  | II | 9 | 9 | 8 | 8 | 8 | 8 |
|  |  | III | 7 | 8 | 8 | 7 | 7 | 7 |
|  |  | IV | 25 | 25 | 24 | 24 | 25 | 25 |
|  |  | V | 21 | 23 | 19 | 20 | 21 | 20 |
| Missing | Illumina | PE | 51 | 21 | 16 | 20 | 18 | 20 |
|  |  | PPE | 12 | 6 | 3 | 4 | 5 | 3 |
|  | H37Rv | PE | 7 | 8 | 7 | 7 | 7 | 7 |
|  |  | PPE | 6 | 8 | 6 | 7 | 7 | 7 |
| Overlap | L2 Repeat | PE | - | - | - | 5 | 0 | 0 |
|  |  | PPE | - | - | - | 11 | 5 | 5 |
| Specificity | Lineage | PE | 4 |  |  | 3 |  |  |
|  |  | PPE | 3 |  |  | 0 |  |  |
| Pseudo | Yes | PE | 6 | 17 | 9 | 6 | 7 | 4 |
|  |  | PPE | 4 | 1 | 4 | 9 | 7 | 8 |

### Protein extraction and mass spectrometry

Proteomics samples analyzed by MS in this study were those prepared from the six clinical reference strains in a previous study (including extraction, digestion, and cleanup, stored at -80° C)^8^. Data-dependent acquisition PASEF (ddaPASEF) data were newly acquired using a TimsTOF Pro (Bruker) QTOF mass spectrometer enabled with trapped ion mobility separations connected online to a nanoElute (Bruker) nanoflow liquid chromatography system via a CaptiveSpray source. Solvent composition at the two channels was 0.1% formic acid for channel A and 0.1% formic acid, 99.9% acetonitrile for channel B. 200 ng of peptides were directly injected onto a Picofrit column (75 µm x 450 mm - New Objective, packed with Reprosil C18 Aq, 1.9um, 100Å, Dr. Maisch) maintained at 60° C and separated at 400 nL/min using a gradient from 2 to 27% B in 100 min. We performed ddaPASEF essentially as described with 10 PASEF scans per topN acquisition cycle^31^. Singly charged precursors were excluded by their position in the m/z–ion mobility plane, and precursors that reached a target value of 20,000 arbitrary units were dynamically excluded for 0.4 min. The quadrupole isolation width was set to 2 m/z for m/z < 700 and to 3 m/z for m/z > 700. TIMS elution voltages were calibrated linearly to obtain the reduced ion mobility coefficients (1/K0) using three Agilent ESI-L Tuning Mix ions (m/z 622, 922 and 1,222).

### Generation of iPtgxDBs for the identification of novel small proteins

For each strain, a standard integrated proteogenomics database (iPtgxDB) was created as described^23^. In brief, NCBI’s PGAP annotation (v4.8), *ab initio* gene predictions from Prodigal (v.2.6.3)^39^ and ChemGenome (v.2.1; with parameters: method: Swissprot space; length threshold: 70 nt; initiation codons: ATG, CTG, TTG, GTG)^40^, and *in silico* ORFs (≥ 18 aa) based on a modified six frame translation that considers the three most common alternative start codons (TTG, GTG and CTG), were hierarchically integrated. Minimal redundancy of protein sequences in the iPtgxDB was achieved by joining different versions of a gene with an identical stop codon into annotation clusters. Such clusters include a full-length anchor protein sequence plus different start sites, provided that these longer or shorter proteins (compared to the canonical RefSeq annotation) are detectable by tryptic peptides.The pre-processing step determines the ambiguity level of each peptide with respect to all protein sequences and -as a unique feature-to the encoding gene models (for detail, see https://iptgxdb.expasy.org). Allowing trypsin cleavage before P was enabled and pseudogenes were only included if their stop matched a prediction from another annotation source to prevent cases where peptides would be discarded as ambiguous due to overlapping predictions with different stops, which occur frequently for pseudogenes. We also created smaller custom iPtgxDBs for the 6 strains, (less than 9,800 proteins, i.e., 12 times smaller than the standard iPtgxDBs), which exhibit better search statistics^41^. For these, we replaced the large number of *in silico* predictions with ∼1,500 orthologs of proteins from strain H37Rv (6aa and longer, i.e., detectable by MS) from overall 2,299 pervasively translated CDS identified by ribosome profiling^42^, a valuable data set that may include potential novel SEPs.

### Proteomic data analysis

For each strain, ddaPASEF data in Bruker *.d format (biological triplicates) was searched against a RefSeq protein search DB and the standard and custom iPtgxDBs. All DBs were merged with contaminants from the CrapOme dataset and searched using FragPipe 22.0 with default strict trypsin settings: Peptides were identified using MSFragger v.4.1^43^ and rescored first with MSBooster 1.2.31 using the Prosit_2019_rt and Prosit_2023_intensity_timsTOF models for retention time and peak intensity prediction, respectively, followed by processing with Percolator v.3.6.5 using the –no-terminate and –post-processing-tdc flags. Proteins were inferred using ProteinProphet with default settings and the filter command of Philosopher 5.1.1 was used to obtain an FDR of 0.01 at the PSM, peptide and protein level based on the sequential and picked FDR approach with razor peptides enabled. The results of 3 replicates were merged using Abacus, again with razor peptides and picked FDR enabled. The proteotypicity of all identified peptides was assessed using PeptideClassifier^44^ further extended to support proteogenomics in prokaryotes^23^. Proteins only identified by ambiguous peptides that match to multiple proteins encoded by different gene models (3b peptides) were filtered out.

To minimize spurious hits, an additional ad hoc filter was applied to RefSeq and novel proteins. For RefSeq proteins, at least 2 PSMs were required and at least two peptides to identify proteins above 15 kDa^45^. The RefSeq DB search results are shown in **Supplementary Table S7**. To increase the stringency for novel protein identifications as recommended^29,46^, an annotation resource-dependent threshold of PSMs was applied on top^23^: Novel proteins and start sites of annotated proteins predicted by Prodigal, ChemGenome or orthologs of H37Rv Ribo-seq identifications were required to have at least 3 PSMs, while *in silico* ORFs were required to have at least 4 PSMs. In cases where multiple extensions were detected for the same protein cluster, these entries were collapsed and their PSMs and peptides summed up. Since iPtgxDBs only contain pseudogenes if other predictions with matching stops exist, novel proteins were reclassified as pseudogene identifications if they had an in-frame overlap with an annotated pseudogene that accounted for at least 50% of the length of the shorter of the two sequences.

### Entrapment analysis for novel protein identifications

The recently reported entrapment approach^32^ was utilized to assess the accuracy of the protein-level FDR reported by Philosopher. For this, protein-level entrapment DBs based on the sequences in the iPtgxDBs were created with FDRBench 0.4.0. These were then searched as described, except that a newer version of Philosopher (5.1.2) was used which recently introduced support for protein-level q-values. To obtain false discovery proportion (FDP) values for the full FDR range, all FDR filtering steps of Philosopher (PSM, peptide and protein level), as well as filtering by peptide and protein probability were disabled. As in the normal searches, Philosopher’s Abacus was used to integrate results from multiple samples, but it was extended to calculate strain-level q-values. FDRBench was then used to estimate the global FDP of all detected iPtgxDB entries and then separately for the subset of detected canonical Refseq proteins and the subset of novel iPtgxDB proteins (novel proteins, novel starts and pseudogenes) by only considering the identified target, decoy and entrapment proteins of the respective subsets. The subset-specific error rates were then assessed by visualizing the correlation between the global protein FDR (q-value) and the estimates of the subset-specific FDP using a Python script (see **Data Availability**). For this, the values of all six strains were aggregated into bins of 1% FDR, ploting curves through the means and showing the standard deviation as lighter bands. Sets of high confidence novel proteins were selected by first setting a strain-specific FDR threshold corresponding to a local FDP of 1% among novel proteins and subsequently applying the ad hoc filter, which resulted in much lower protein-level FDR thresholds than the standard 1% (**Supplementary Table S9**).

### Bioinformatic analysis of novel small proteins

#### Analysis of transcriptomics data

RNA-Seq data of the six Mtb strains from^8^ were used to explore Pearson’s correlation for all genes (including novels) that were expressed at the transcript and protein level (**Supplementary Fig. S12**). For more detail, see **Supplementary Methods**.

#### Functional predictions

Novel proteins were annotated based on the eggNOG v5.0 DB using eggNOG-Mapper v.2.1.12^47^ in diamond mode and with Phyre2^48^ using default options. The subcellular localization and presence of signal peptides and other motifs was predicted using InterProScan v5.59-91.0^49^, extracting predictions from TMHMM, SignalP and Phobius, and with the command line version of PSORTb 3.0^50^ (using docker image v1.0.2) and LipoP 1.0a. Potential operons were predicted using OperonMapper^51^ for the combined list of RefSeq genes and novel protein-coding ORFs (excluding pseudogenes). Potential antimicrobial peptides (AMPs) were predicted with AMP Scanner (v2)^52^, as described^28^. A distribution of prediction scores was created for all RefSeq and novel proteins; novel CDS below 200aa with a probability score greater than 0.99 were considered potential AMP candidates with higher confidence.

#### Conservation

The sequences of novel protein candidates were searched against the NCBI nt DB using tblastn 2.12.0 (word size 6). The number of hits (requiring >= 60% sequence coverage, >= 40% sequence identity and an e-value < 0.01) was assessed at different taxonomic ranks using the ete3 python library (v.3.1.2). Taxonomic ranks were marked if at least 10 additional hits were observed^41^.

#### Signal of purifying selection

Novel CDS candidates were analyzed for a signal of purifying selection using the same criteria as described^42^: The start and stop codons, as well as codons overlapping with annotated genes were removed from the nucleotide sequences to prevent biases in the analysis (several novel proteins completely overlapping with an annotated gene were thus excluded). The presence of a significant difference in the G/C frequency (ratio of G and C to A and T) between the second and third positions of the codons was checked for each candidate using Fisher’s exact test. Novels were regarded as being under purifying selection if the P-value of the Fisher’s exact test was below 0.1 and a positive G/C frequency ratio between the third and second codon position existed, i.e., a higher G/C frequency at the 3rd codon (**Fig. 5G**). The conservation of these novels across several taxonomic ranks including Mycobacteriaceae was also assessed.

#### Pangenome analysis

The lists of annotated and novel protein-coding genes (excluding pseudogenes) were compared using Panaroo 1.5.2 to determine whether novel proteins were found in only one, a subset (e.g. lineage specific) or multiple strains. Predictions for the novel proteins were merged per orthogroup, keeping unique values which arise due to sequence differences of the orthologs between strains. The only exceptions were the number of PSMs and identified peptides, which were kept separate per strain, and the conservation BLAST results, for which the maximum value among the strains was used at each taxonomic level. To identify novel proteins that are orthologous to actively translated mRNAs identified by Ribo-seq in H37Rv^42^, a second pangenome analysis was run with Panaroo using annotated and Ribo-seq identified protein-coding genes of H37Rv as additional set.

#### Data Availability

Genome sequences of the six Mtb strains (BioProject PRJNA544196, BioSamples SAMN11822579-84) are available from NCBI Genbank with the accession numbers (acc#) CP040688.1 (N0052), CP040689.1 (N0072), CP040690.1 (N0153), CP040691.1 (N0155), CP040692.1 (N0157) and CP040693.1 (N0145). Illumina (I) read data from^6^ and PacBio (PB) read data from^7^ were used; they are available from the European Nucleotide archive: N0052 (acc# for I: ERR2704677, ERR2704699, ERR2704698; for PB: ERR956959, ERR964405), N0072 (acc# for I: ERR2704680; for PB: ERR956956, ERR964402), N0145 (acc# for I: ERR2704702, ERR2704701, ERR2704683; for PB: ERR956958, ERR964404), NO153 (acc# for I: ERR3183990; for PB: ERR956957, ERR964403), N0155 (acc# for I: ERR2704703, ERR2704684; for PB: ERR964407 & ERR964414), N0157 (acc# for I: ERR2704704, ERR2704685; for PB: ERR956955 & ERR964401). RNA-Seq data from^8^ are available on ArrayExpress with the dataset identifier E-MTAB-8103. The timsTOF proteomics data will be available at the ProteomeXchange Consortium via the PRIDE partner repository (acc#: PXD081163, reviewer token: aW7asyHcUcUx). The Snakemake pipeline for the classification of PE/PPE genes is available on GitHub (https://github.com/mgbteam/mtb_pe_ppe), as is the code for the comparative genomics and stringent proteogenomics analyses (https://github.com/mgbteam/mtb_paper).

## Results

### Meta-analysis of *M. tuberculosis* genome assemblies

By the end of 2024, overall 7,193 Mtb genome assemblies have been deposited at NCBI RefSeq, only 429 of which (6.0%) fell into the highest quality category “complete genome”. The others belonged to the lower assembly quality level categories “chromosome” (256), “scaffold” (4,243), or “contig” (2,265) (**Fig. 1A**). Based on available metadata, 326 assemblies (4.5%) were generated using long read sequencing data (PacBio, ONT) or from a hybrid approach (combining long and short reads). In contrast, 4,703 were based on short read sequencing data (Illumina or other), while no information was made available for 2,164 assemblies (**Supplementary Fig. S1**). Assemblies in the categories scaffold and contig comprise a larger number of contigs. Based on the median of 66 contigs observed for all Mtb assemblies (**Fig. 1B**), we can safely assume that the vast majority of assemblies in the category “unknown” are based on short read data. This is relevant, as pangenome estimates based on short read datasets can be artificially inflated and lead to incorrect assumptions about evolutionary scenarios^21^.

**Fig. 1.**
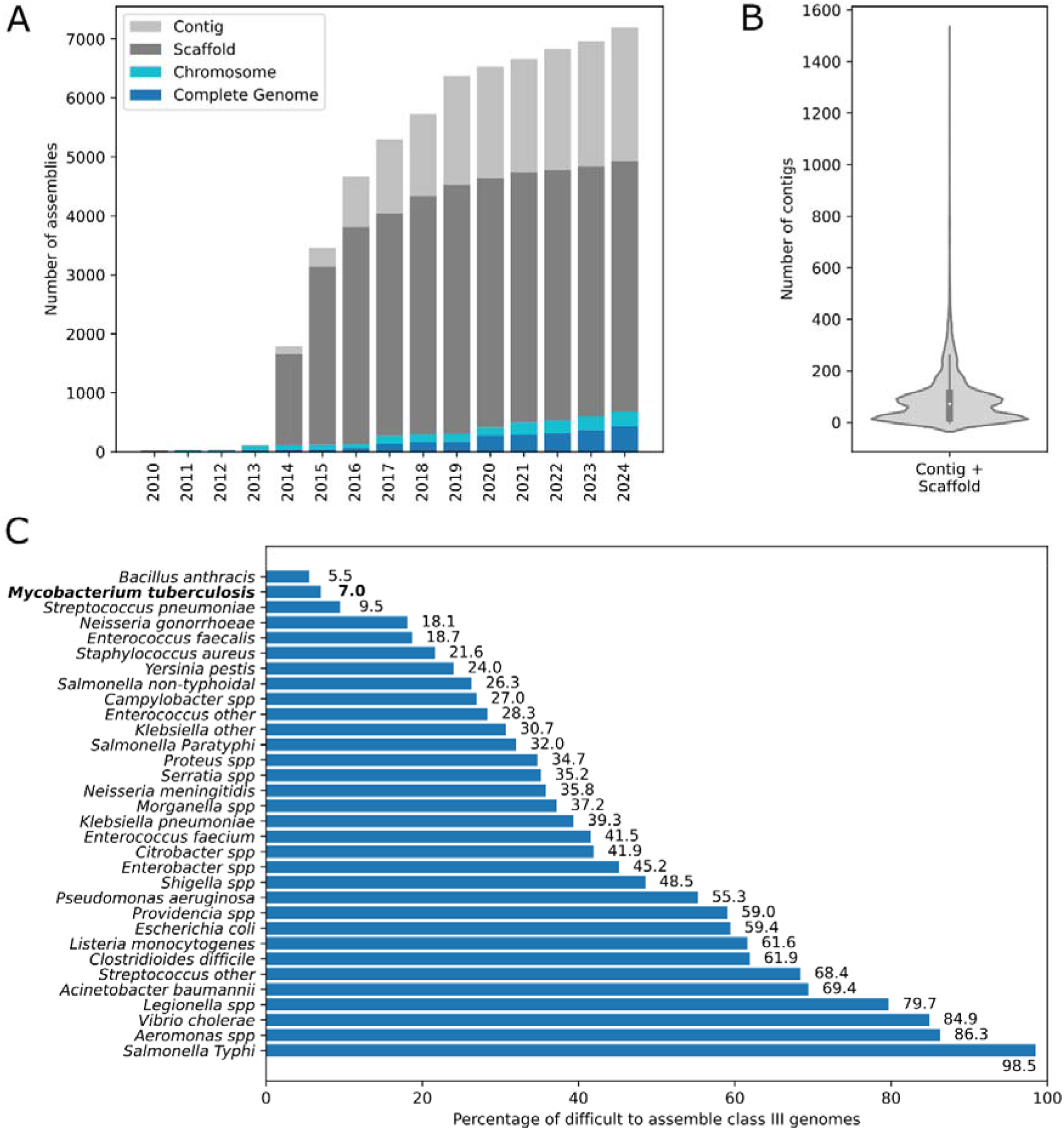
Meta-analysis of genome assemblies of *M. tuberculosis* & major pathogens. **A**. Increase of publicly available NCBI RefSeq genome assemblies for *M. tuberculosis* from 2010 to 2024 colored according to NCBI’s four assembly quality categories: complete genome (dark blue), chromosome level (light blue), scaffold (dark gray) and contig (light gray). **B**. Violin plot of the number of contigs reported for the roughly 6,500 RefSeq *M. tuberculosis* assemblies in the categories scaffold and contig. **C**. Genome assembly complexity, here shown as percentage of difficult to assemble class III genomes, for a selection of major human bacterial pathogens (see **Supplementary Fig. 1B**). *M. tuberculosis* is shown in bold.

We also analyzed the genome assembly complexity of the 429 Mtb strains with complete genomes (see **Methods**): 399 genomes (93.0%) represented easy to assemble class I genomes. While no repeat-rich class II genomes were found, 30 genomes (7.0%) were difficult to assemble class III genomes, with repeats longer than the rDNA operon. Such repeats can extend well over 100 kb in length^19^, which has been described for L2 strains of Mtb^53^. Next, we compared the assembly complexity of over 30 bacterial pathogens, including species in the list of critical and high priority pathogens from the WHO 2024 report^70^ and species or genera from the Global Burden of Disease Study^34^ (**Supplementary Fig. S1B**). Strikingly, Mtb has the overall second lowest percentage of difficult to assemble class III genomes (7.0%), after *Bacillus anthracis* (5.5%) (**Fig. 1C**). Mtb is thus very well suited for genomic and proteogenomic investigations.

### *De novo* genome assembly of six clinical Mtb reference strains

To create an optimal basis for comparative genomics and proteogenomics including novel small protein discovery, we *de novo* assembled the genomes of three clinical reference strains of L1 (N0072, N0153, N0157) and L2 (N0052, N0145, N0155), respectively. They were selected from the set of 20 clinical reference strains described above^6^ (**Fig. 2A**), and for which Illumina data was available. Long PacBio reads from a later study^7^ were assembled with the assembly algorithm Flye before polishing with short Illumina reads (**Supplementary Methods**), thereby reaching a combined read coverage of at least 200-fold (**Supplementary Table S2**). Each strain harbored a single chromosome of 4.40 to 4.42 mega base pairs (Mb) with a GC content of 64.7 to 65.2% without additional plasmids. The coverage plots of the PacBio reads indicated that while the three L1 strains were completely resolved, all three L2 strains contained an unresolved repeat region of around 240-380 kb in length with an up to two-fold higher coverage (**Fig. 2B**, **Supplementary Fig. S2**; **Supplementary Table S2**). Such large duplicated regions have been reported for a subset of Mtb strains of the W/Beijing family of L2^53^. Repeats of this size are currently not resolvable without very substantial additional effort like optical mapping. For the L2 strains, we thus report their genomic location, estimated repeat region size (**Supplementary Table S2**) and annotated genes contained (**Supplementary Table S3**).

**Fig. 2.**
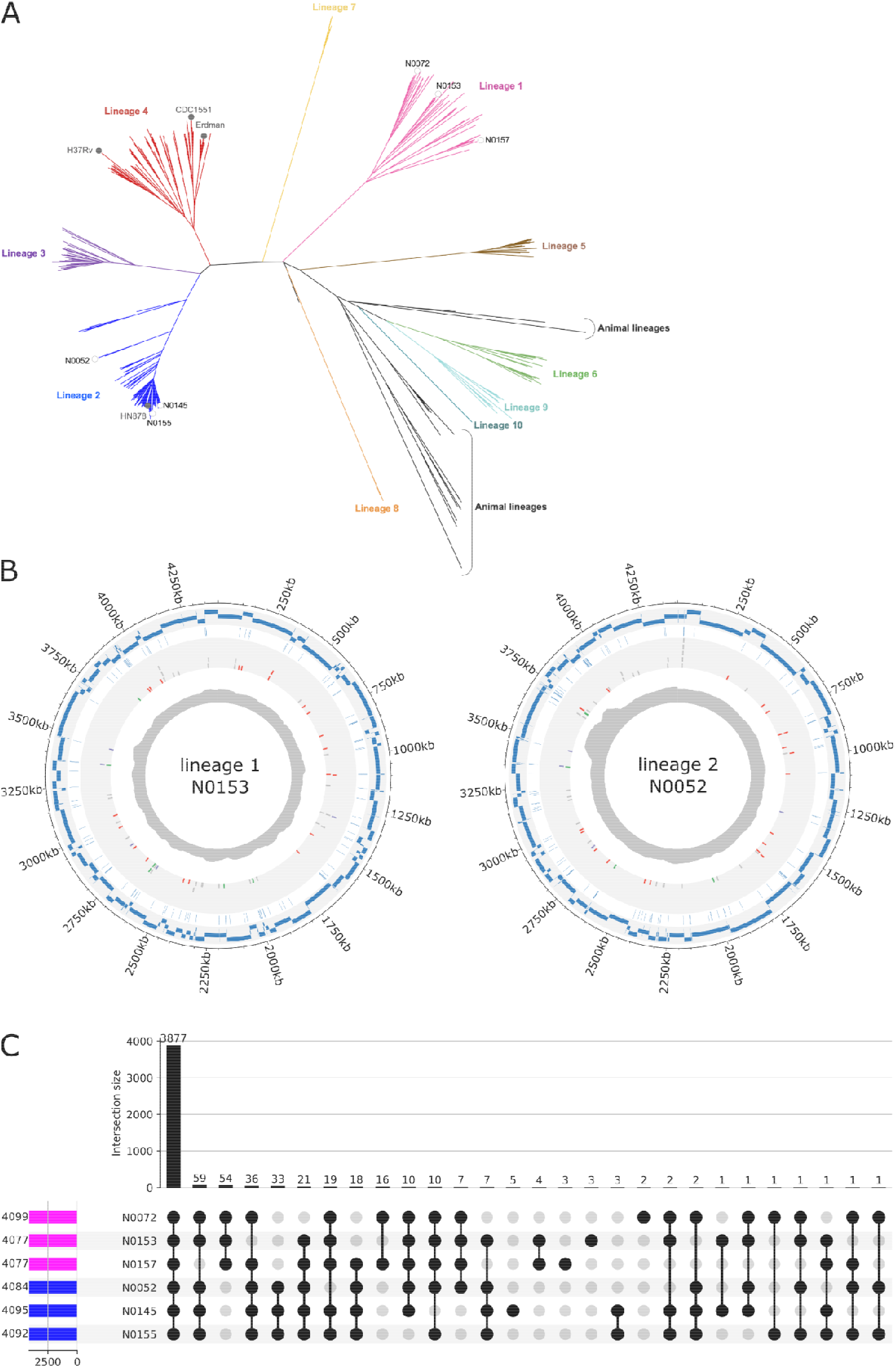
Complete genomes of six clinical reference strains of L1 and L2. **A.** Maximum likelihood topology of the six L1 and L2 clinical strains (open circles) analysed here plus 278 genomes representative of MTBC global diversity (10 lineages). Gray dots indicate the phylogenetic placement of four widely used *M*. *tuberculosis* laboratory strains. Branch lengths are proportional to nucleotide substitutions; adapted from^6^. **B.** Comparison of our complete *de novo* assemblies to fragmented Illumina assemblies for strain N0153 of L1 and N0052 of L2. The outer rings show the complete genome, followed by the Illumina contigs (blue bars), the observed gaps (blue dashes), and a circle for genes partially missed/incomplete (gray dashes) where three enriched gene families are shown: PE genes (red dashes), PPE genes (green dashes) and transposons (purple dashes). The PacBio coverage is shown in the innermost circle. **C**. Upset plot showing the number of core genes and those conserved in various subsets. These include lineage specific genes, i.e., genes that are conserved and specific either for L1 (54) or L2 (33).

Next, we compared our complete and contiguous PacBio genomes against assemblies only based on Illumina short read data^6^. One example from L1 and L2 is shown in **Fig. 2B**, the other four in **Supplementary Fig. S3**. While the short reads covered ∼99% of the genome sequence when considering multiple mapping reads (except for strain N0072 with 97% coverage), they partially or completely lacked between 43 to 160 CDS (**Supplementary Table S1**), i.e., roughly 1-4 % of the annotated CDS. Such genomic “blind spots” can contain highly relevant genes (see Introduction), as shown for *P. aeruginosa*^11^. Among the CDSs not completely covered in the Illumina assemblies, we observed an enrichment for transposases or insertion sequences (P-value <0.01 except for strain N0155) and a highly significant enrichment for PE or PPE family members with ∼20 or more members missing per strain (all P-values < 1.05E-16) (**Supplementary Table S4**).

### Comparative genomics

A comparative genomics analysis of the six strains using Panaroo^38^ identified 4,198 orthologous groups (orthogroups) overall, with a large core genome of 3,877 genes (92.4%) (**Fig. 2C**, **Supplementary Table S5**). 308 genes (7.3%) were classified as accessory, i.e., present in a subset (two to five) but not in all strains, and 13 genes as strain-specific, i.e., occurring only in one strain. The strain-specific genes include 8 hypothetical proteins, a DUF222 domain containing protein, 2 PPE family proteins (including a second copy of PPE38 unique to strain N0153) and 2 type VII secretion target proteins (WXG 100 family^54^). Among 87 genes conserved in all three strains of one lineage, 54 genes (1.3% of all genes) were specific for L1 and 33 genes (0.8%) specific for L2 (**Fig. 2C; Supplementary Table S5**). L1-specific genes included 7 PE/ PPE family members, multiple CRISPR-associated proteins (type III CRISPR proteins Csm4-6, Cas1 and Cas2 endonucleases), a type II toxin-antitoxin system VapB family antitoxin and several proteins involved in transport (an RND family transporter, a mechanosensitive ion channel, a cation-transporting P-type ATPase, an FtsX-like permease family protein and the transport accessory protein MmpS). Conversely, L2-specific genes included the manganese-binding transcriptional regulator MntR and a LuxR family transcriptional regulator, the gene encoding sensor histidine kinase KdpD, part of a two-component system for which a role in virulence and intracellular survival of pathogenic bacteria including mycobacteria has been described^55^, two enzymes with roles in triacylglycerol metabolism, which has been suggested as an energy source for the persisting, slow metabolizing Mtb population in the host^56^ and 3 PE family members (**Supplementary Table S5**).

#### Complete catalogs of PE and PPE genes

PE and PPE genes exhibit a high GC content and extensive repetitive homologous sequences, which has impeded their molecular and biochemical analysis^13,14^ and explains their enrichment in genomic blind spots of short read-based assemblies. The comparative genomics analysis identified 4 PE and 3 PPE genes specific for L1 and 3 PE genes for L2 (**Table 1**, **Supplementary Fig. S4**). Due to their relevance, we classified the sets of PE and PPE genes from all six strains as predicted by PGAP, InterPro and orthologs from strain H37Rv into sub-families based on the presence of specific amino acid motifs and a detailed classification of 99 PE and 69 PPE genes of H37Rv^12^. We also assigned sub-lineages based on a phylogenetic analysis of the conserved N-termini (95 aa for PE, 180 aa for PPE family members) as reported^13^ (see **Methods**, **Supplementary Methods**, **Table 1**, **Supplementary Table S6**). Per strain, 87 - 97 PE genes and 60 - 66 PPE genes were identified, plus an additional 4-17 PE and 1-9 PPE pseudogenes (**Table 1**). Most PE genes were assigned to sublineage 5, which contains the PGRS subfamily, while for PPE genes the sublineages IV and V (and MPTR and SVP subfamilies) were most frequent.

A pangenome analysis of the six strains plus H37Rv identified 13 to 16 PE/PPE genes in clinical strains, but not in H37Rv (**Table 1** and **Supplementary Table S6**). Notably, 5 to 16

PE/PPE genes were located in the large repeats of the L2 strains (**Table 1**), and all PE genes missed in Illumina assemblies were PGRS subfamily members. A detailed list of identified PE/PPE genes, including pseudogenes, lineage-specific and clinical reference strain-specific members is provided in **Supplementary Table S6,** the configuration of the *ppe38-71* gene locus in **Supplementary Fig. S5** and detail from the original classification^12^ into subfamilies and sublineages in the **Supplementary Material**. A Snakemake pipeline for this comprehensive identification and classification for other Mtb isolates is made publicly available (see **Data Availability**).

The repetitive sequences of PE and PPE genes also impair their detectability by MS. A PeptideClassifier^44^ analysis identified i) a 2-fold (PE) to 5-fold (PPE) higher percentage of ambiguous peptides (**Supplementary Fig. S6**), and ii) that at least one unambiguous MS-detectable peptide (6 to 40 aa) existed for 92 to 96% of all PE/PPE genes per strain. At least in theory, most gene products of both families can thus be uniquely identified. All fully tryptic unambiguous peptides are reported in **Supplementary Table S6**.

### Proteomic analysis of the six clinical Mtb reference strains

To establish a baseline, we first analyzed the MS proteomics data against a RefSeq-derived canonical set of protein sequences. We measured unfractionated total cell extracts with the timsTOF Pro mass spectrometer using ddaPASEF^31^, a mode that allows to fragment more MS1 precursors and increases the number of acquired MS/MS spectra. We also relied on computational tools to increase the number of correct peptide spectrum matches (PSMs), i.e., MSBooster. Around 2,700 to 2,790 of roughly 4,240 RefSeq proteins per strain were identified at a protein FDR well below 1%, i.e., a coverage of roughly 66-68% of the respective theoretical proteome (**Supplementary Table S7**). This included 167-186 annotated short proteins ≤ 100 aa, i.e., 31-35% of the theoretical small proteome. Products of several CDS that were partially or entirely missed in the Illumina assemblies of L1 and L2 strains were identified in the RefSeq search (**Supplementary Table S8**). These included the small ESAT-6-like proteins EsxL (94 aa) in 5 of the 6 strains and EsxN (94 aa) in 2 of the 6 strains (related proteins encoded by distinct genes, for which some peptides match both proteins (class 3b), while others are unambiguous class 1a peptides), members of the WXG100 family of secreted or secretory-type proteins, and the type VII secretion system ESX-1 associated protein EspI in 4/6 strains. Four PPE family genes missed in the more fragmented N0072 assembly were also identified, and a partially missed PPE gene in N0145.

### Proteogenomics identifies missed protein coding genes in sets of phylogenomically related Mtb clinical strains

We next searched the ddaPASEF data against strain-specific iPtgxDBs. By integrating annotations from reference genome centers and additional predictions (**Fig. 3**, panel “Strain-specific proteomics search databases”), these databases capture almost the entire coding potential of prokaryotic genomes and allow to identify novel CDS^23^. A pre-processing step (**see Methods**) ensures that around 95% of all peptides uniquely identify one protein (see https://iptgxdb.expasy.org/) despite the much bigger DB size (∼69,000 gene clusters and 121,000 proteins in a standard iPtgxDB compared to ∼4,240 proteins in a RefSeq DB, **Supplementary Table S11**). Notably, for Mtb, we could rely on a Ribo-Seq dataset of actively translated mRNAs from H37Rv^42^, which could include missed CDS. We replaced the *in silico* ORFs with ∼1,500 orthologs in the size range detectable by MS to create additional, much smaller, custom iPtgxDB (∼9,800 proteins, **Fig. 3**) for each strain. While the upstream genomics (check if complete genomes exist, otherwise de novo assemble and annotate; **Fig. 3**, right upper panels) and proteomics aspects (extract proteins, digest and analyze by tandem MS) of our proteogenomics workflow were executed with minor adaptations - meta-analysis for Mtb genomes, ddaPASEF for tandem MS-, we substantially adapted the downstream part for the analysis of phylogenomically related strains (two bottom rows of panels, **Fig. 3**): we added a stringent FDR control by entrapment (see below) and integrated the comparative genomics results of the six strains (**Fig. 3**, panel “Pangenome of strains”), both for annotated and newly identified protein coding genes (again computed with Panaroo). We show them as groups of novel genes, of novel start sites whose annotations differ from RefSeq and of expressed pseudogenes, i.e., the three types of novel information that can be uncovered by proteogenomics (**Fig. 3**, lowest panel). We refer to these conserved groups as orthogroups from here on. The search against the standard iPtgxDBs identified 2,949-,3060 proteins per strain that were identified with unique peptides at a protein FDR below 1% (Philosopher). To limit the number of potential false positive identifications, we applied an additional resource-specific PSM and peptide filtering as recommended^29,46^. We require more spectral evidence from ab initio gene predictions and Ribo-Seq orthologs (≥ 3 PSMs), from *in silico* predicted ORFs (≥ 4 PSMs) and at least 2 peptides for longer proteins (≥15kDa). 2,729-2,824 proteins per strain remained after this “ad hoc” filter (**Table 2**). Overall, unambiguous peptide evidence was observed for all three types of novelty (**Fig. 3**), including 12-24 novel proteins per strain (with up to 3 novel CDS uniquely contributed by the custom iPtgxDB), 56-60 longer and shorter forms of RefSeq annotated CDS (**Table 2**), and 8-17 genes annotated as pseudogenes. Notably, the novel proteins were enriched in SEPs (54-75%, **Table 2**).

**Fig. 3.**
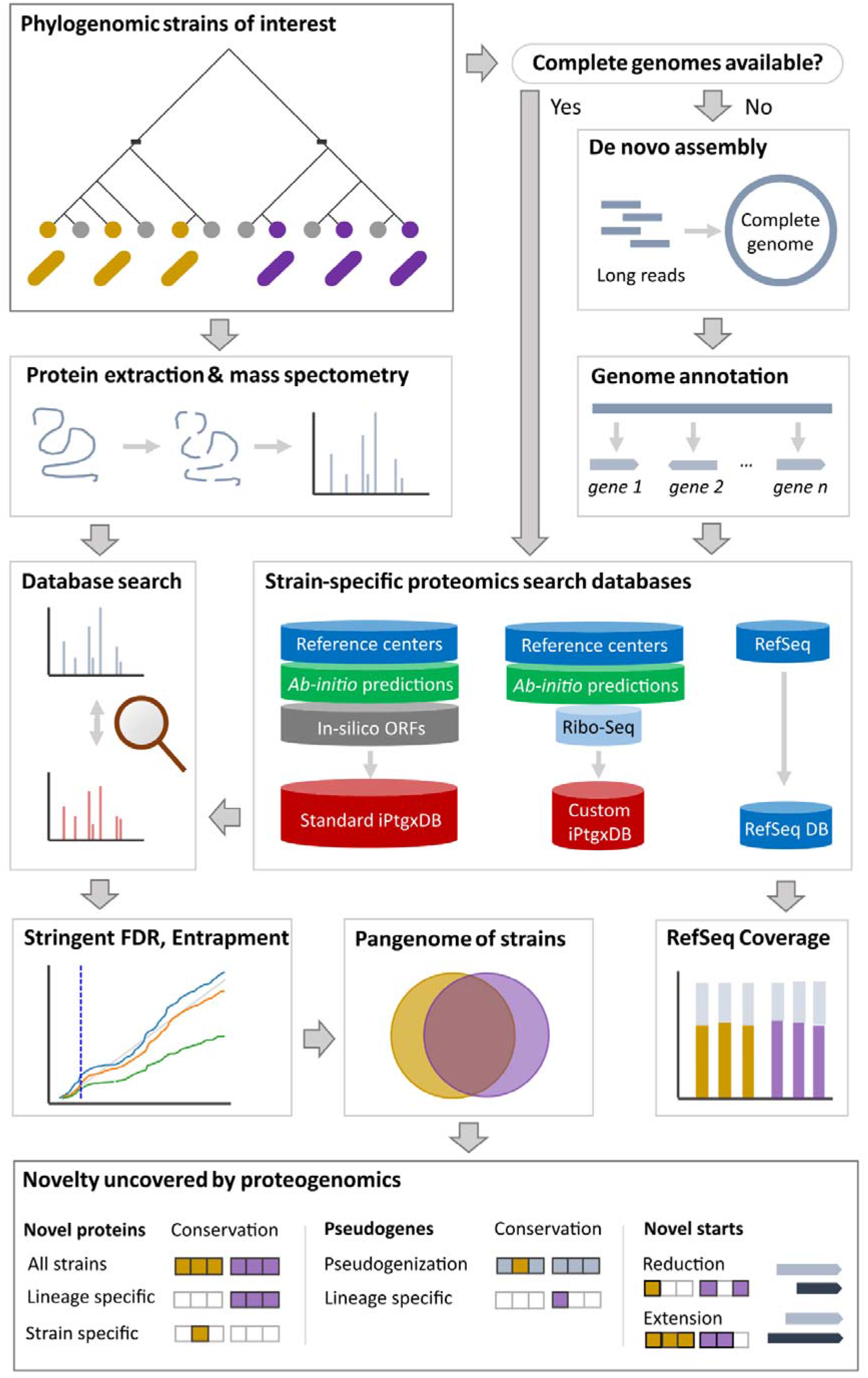
A generic approach for proteogenomics on sets of related strains. Phylogenomically relevant sets of related strains that ideally capture the diversity within and between lineages are selected. If no complete genomes are available the strains are sequenced with long reads, assembled de novo and annotated (right upper panels). From the strains, protein extracts are generated and analyzed with a suitable proteomics workflow (panel “Protein extraction and mass spectrometry”), in our case ddaPASEF. The acquired tandem mass spectra are searched against a RefSeq DB to deduce the coverage of the theoretical proteome (panel “RefSeq coverage”), and an iPtgxDB (panel “Strain-specific proteomics search databases”) to identify the three types of novelty. Based on an available Ribo-Seq dataset, we also created a much smaller custom iPtgxDB. Entrapment (panel “Stringent FDR, Entrapment”) is used as a very stringent approach to control the local FDR of novel proteins beyond simple filtering based on PSM thresholds. Relying on a pangenome analysis of annotated and novel genes, the downstream analysis is simplified and the data presented in the form of conserved orthogroups over the six cell lines in three classes of novelty.

**Table 2.**
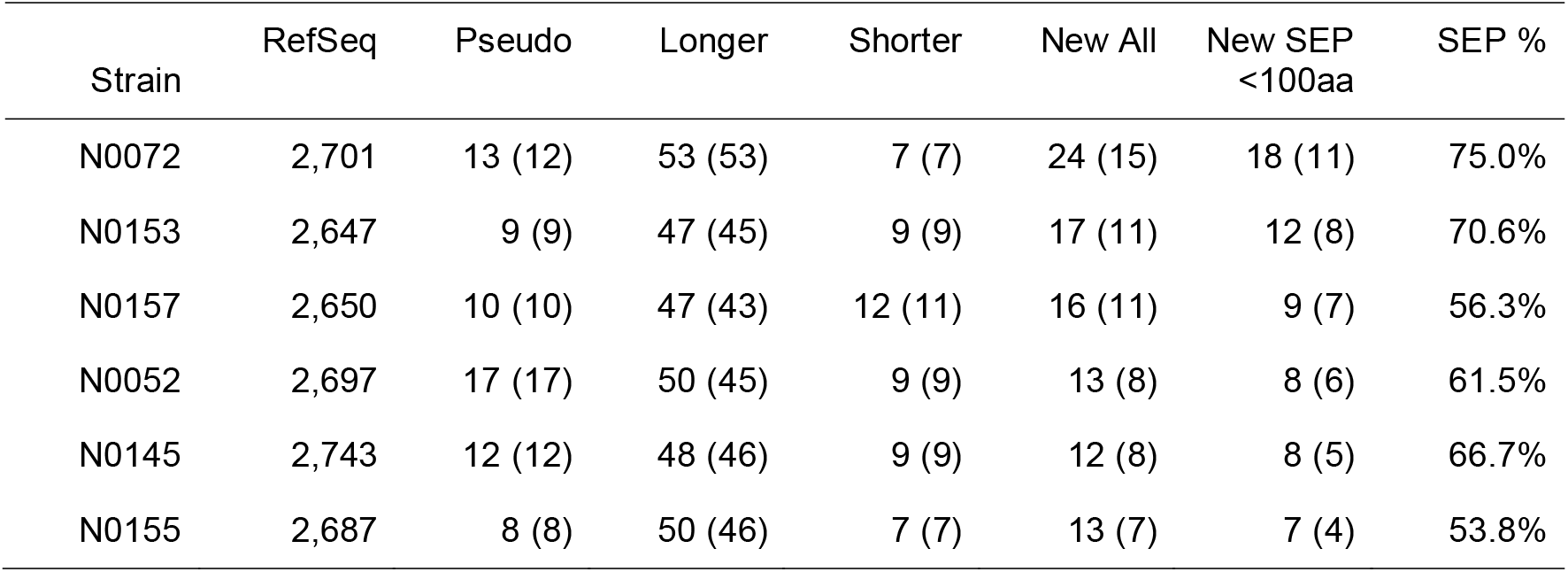
Proteins identified by searches against strain-specific standard iPtgxDBs (5 novels from custom iPtgxDB searches are included in columns New ALL, New SEP). Higher confidence novel identifications that pass an additional protein-level FDRBench filtering step (see next section) are shown in brackets.

### FDR estimation and control of proteogenomic novelty

An important adaptation to our workflow (**Fig. 3**) was to carry out entrapment based estimation of the FDR relying on a recently published approach^32^. To address the well described problem of a much higher FDR among novel proteins compared to kown proteins in proteogenomics^46,71^, we not only calculated the false discovery proportion (FDP) for all detected proteins, but also assessed the FDP separately for the subset of detected canonical RefSeq and novel proteins. FDRBench calculates 3 metrics to estimate the FDP: a lower bound and two estimates of the upper bound (paired and combined FDP). The estimated FDP for all detected proteins at an FDR of 1% was close to 1% (**Fig. 4A**, left panel), and even lower for the subset of RefSeq only proteins (**Supplementary Fig. S9** and **S10**). However, when considering target and decoy hits from the subset of novel proteins (including novel starts and pseudogenes), the estimated local FDP at a global FDR of 1% was significantly higher, with estimates ranging from 11% (lower bound) to 23% (combined FDP) (**Fig. 4A**, center panel). Applying the ad hoc filter reduced the local FDP at a global FDR of 1%, with the new estimates ranging from around 6% (lower bound) to 11% (combined FDP) (**Fig. 4A**, right panel).

**Fig. 4.**
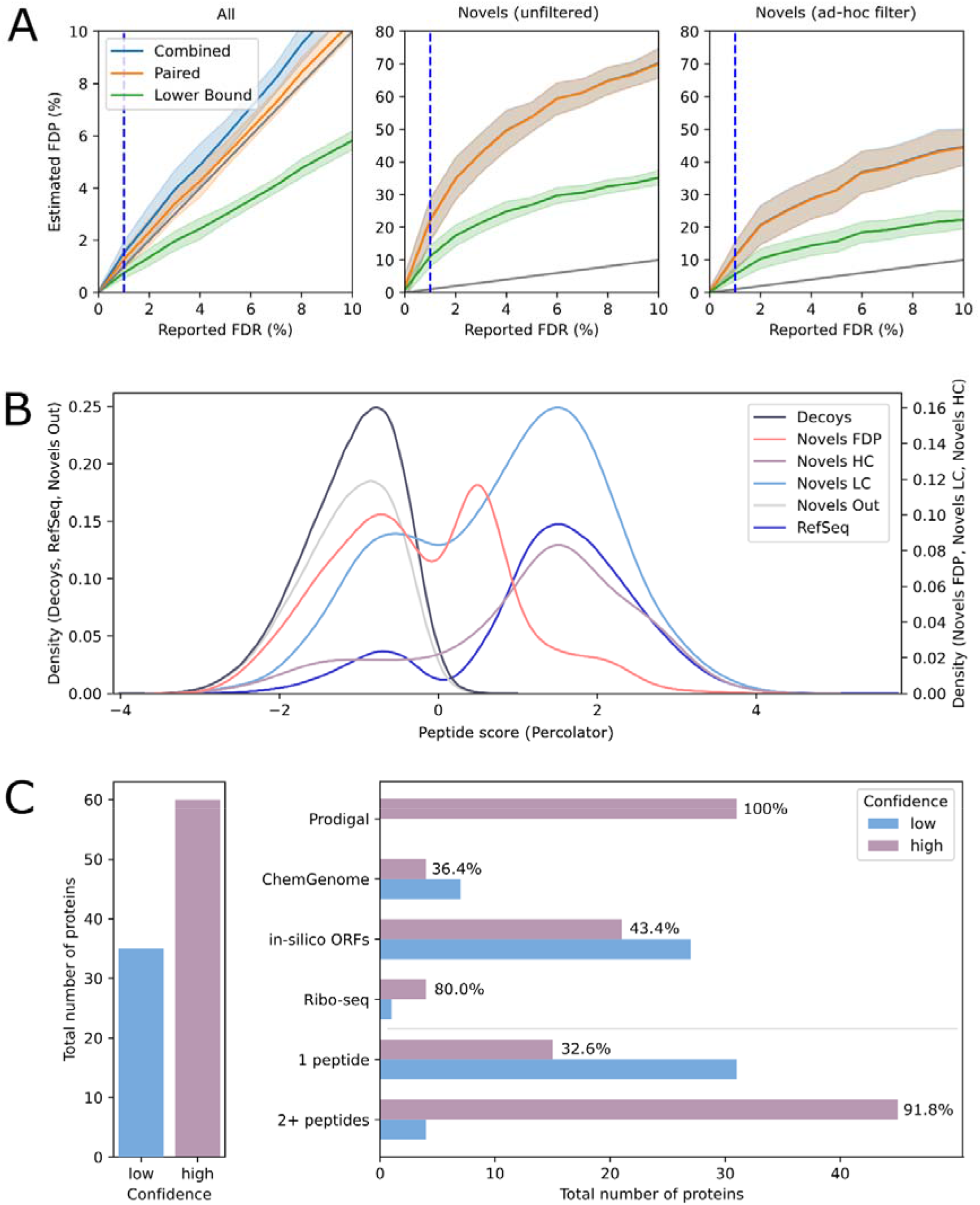
FDR control of proteogenomic novelty **A**. Analysis of the estimated local FDP versus global FDR threshold for i) all identifications from all strains (left panel), and ii) for the subset of novel proteins, pseudogenes and proteins with novel start sites either before (center panel) or after applying the ad hoc filter (right panel). The standard error is shown as lighter bands to show the variation between the six strains. The 1% FDR threshold is shown as a blue dotted line. **B**. Percolator score distribution of peptides for various subsets, summarized over 6 strains with two y-axes for different scales. The score distributions of decoy and filtered out novels are clearly separated from those of RefSeq, high and lower confidence novels. **C**. Percentage of proteins that are classified as higher confidence (HC; purple) when applying the entrapment FDP filter after our ad hoc filters for all novel proteins (left panel) and per annotation source and number of identifying peptides (right panel).

While the ad hoc filter significantly reduced the FDP, it still exceeded the targeted FDR of 1%. We therefore explored applying the FDP estimation as an additional filter. To do so, a new FDR threshold was defined corresponding to an FDP of 1% in the subset of novel proteins **(Supplementary Table S9**). Novel proteins passing this threshold in addition to the ad hoc filter were then classified as higher confidence identifications (see also **Supplementary Fig. S11** and **Table 2**; values in brackets indicate identifications confirmed by FDRBench), those only passing the ad hoc filter as lower confidence (LC). This approach is also supported by analyzing the Percolator score distribution of peptides from different subsets: we noted a clear separation of the score distributions of decoy hits and novels that did not pass either filter (black and gray curves, **Fig. 4B**) from RefSeq proteins and novels passing the ad hoc filter. In contrast, the novels that only passed the FDP filter (red curve) showed a less clear signal and were therefore also discarded. The score distributions of the subset of peptides identifying HC and LC novel hits (light blue and dark blue curves) were similar to that of RefSeq proteins, lending further support to our decision to report both sets.

The FDP-based HC and LC classification impacted the three types of novelty we finally report: summed up over all strains, the large majority of new start sites and expressed pseudogenes (94.8% and 98.6% respectively, **Table 2**) were classified as HC. In contrast, 36.8% of the novel proteins were classified as LC identifications (**Table 2**, **Fig. 4C** left panel). Moreover, we found striking differences for the pass rate for different prediction sources: all hits from the *ab initio* predictor Prodigal were classified as HC, while that rate was substantially lower for ChemGenome (36.4%) and in silico predictions (43.4%). The pass rate among the 5 novels predicted by Ribo-seq detected orthologs in H37Rv was 80%. Importantly, close to 60% of novel identifications identified with 1 peptide over all strains was classified as LC, even if some peptides were identified by many PSMs (up to 42, **Supplementary Table S10**), and in several strains (novel_8 in 4/6 strains, novel_13 in 3/6). In contrast, 91.8% of the identifications with 2 or more peptides were HC (**Fig. 4C**). Notably, all four resources contributed unique novel candidates not covered by other predictions, a benefit of the iPtgxDBs^23^.

### Pan-genomic consolidation and prioritization of types of novelty

A visualization of the pan-genomic orthogroups readily identifies candidates encoded and/or expressed in all six strains, in a specific lineage or in one strain (**Fig. 5A**): the 12-24 novel proteins were collapsed into 37 groups, 32 from searches against the standard iPtgxDBs and five (novel 33 to 37) uniquely identified by searches against the Ribo-seq informed custom iPtgxDBs (**Fig. 5A**, lower panel). Notably, with a median length of 72 and 76 aa, the novel proteins identified with both iPtgxDBs (a Venn diagram shows the overlap of the respective orthogroups identified, **Fig. 5B**) were significantly shorter than proteins from the categories RefSeq (P-values < 3.9 E-6 and < 5.7 E-4, respectively), pseudogenes (Pseudo) and altered RefSeq CDS (New Start), all with a median length of around 300 aa (**Fig. 5B**). The 8-17 expressed pseudogenes per strain were collapsed into 44 orthogroups (**Supplementary Fig. S7**), while the 56-60 novel start sites per strain were collapsed into 70 groups of longer CDS (82%) and 15 groups with evidence for a shorter CDS (**Fig. 5C**, **Supplementary Fig. S8**). Among these 85 orthogroups, a new start site was seen in all six strains in 33 cases (27 extensions, 6 reductions) and in a specific lineage in 22 cases (15 L1, 7 L2; **Supplementary Fig. S8**). For 37 of the 44 pseudogene orthogroups, a matching bona-fide RefSeq CDS was annotated in most other strains; the 7 orthogroups that did not have a matching RefSeq CDS in any other strain were all unique to L2 (**Supplementary Fig. S7**). Notably, 30/37 pseudogenes that were grouped with RefSeq CDS of other strains were annotated as pseudogene in only 1 strain, in line with findings that pseudogenization acts as an important driver of strain diversity in Mtb^57^ which lack HGT.

**Fig. 5.**
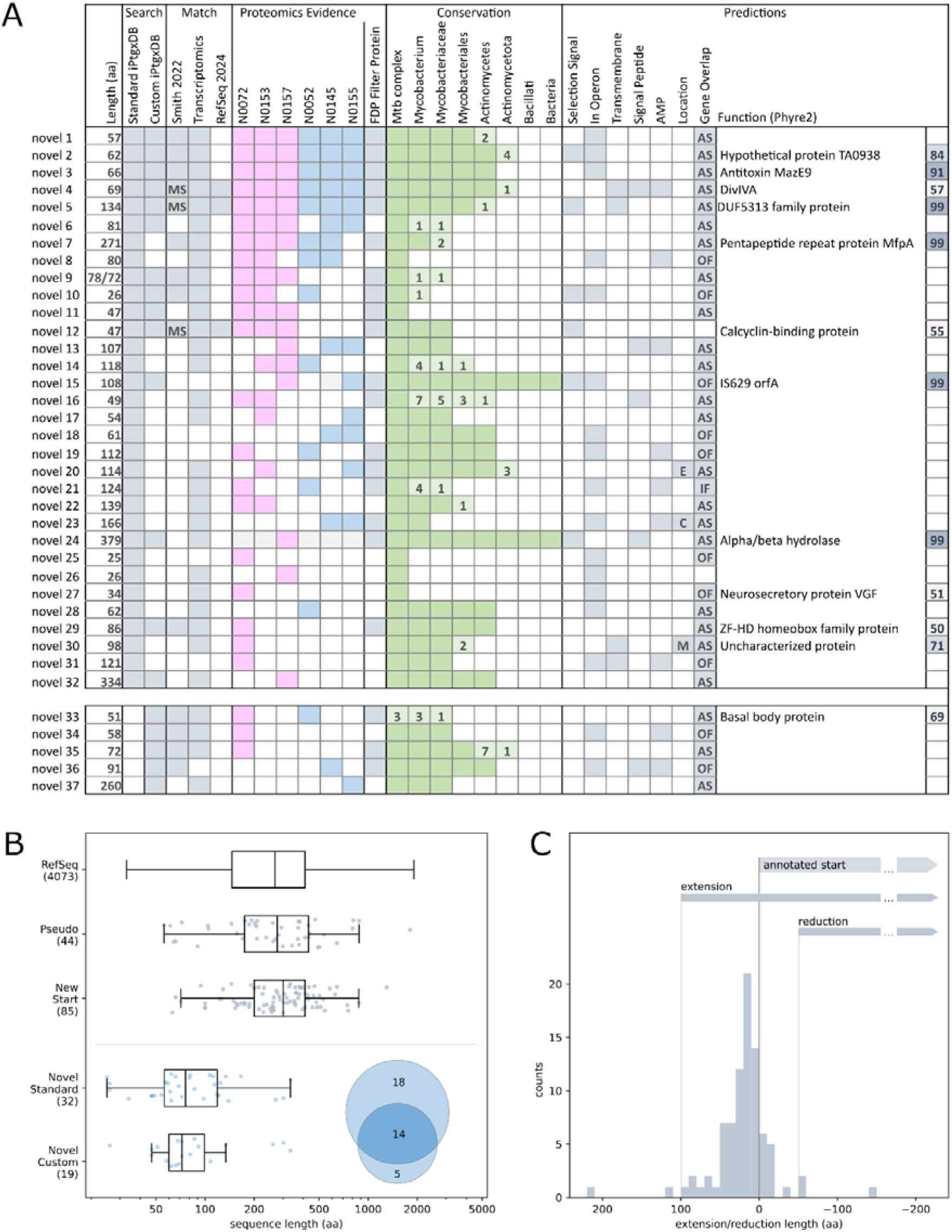
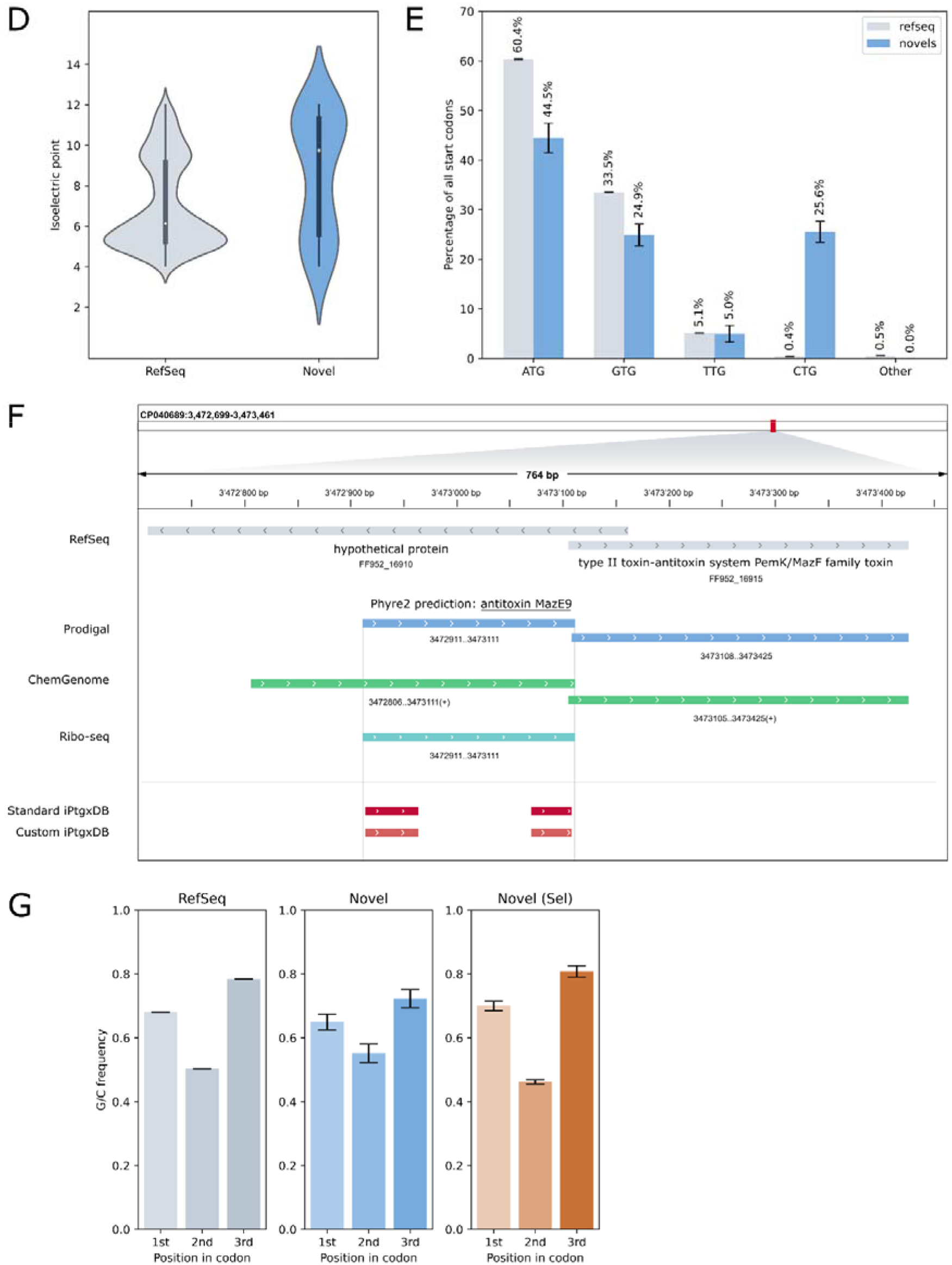
Novel CDS identified by proteogenomics and selected supporting evidence. **A**. Overview of 37 orthogroups of novel proteins identified in the iPtgxDB searches; 5 groups (bottom part) were added by the custom iPtgxDB search. Proteomics evidence is denoted as purple (L1) and blue (L2) boxes. The column FDP filter indicates if a protein passed the stringent FDRbench filtering. Orthologs of CDS identified with Ribo-Seq in H37Rv are shown (MS: additional proteomics evidence). Conservation across selected taxonomic groups is shown as green boxes; fully filled boxes (novel ORF identified in at least 10 additional subtaxa), respective confidence score is above 50%. **B**. Boxplots for protein length in 5 categories (summarized over all strains). A Venn diagram with the overlap of novel proteins identified with both iPtgxDBs (right corner). **C**. Summary over new protein start sites implying a longer (extension) or shorter (reduction) CDS than annotated by RefSeq. A schematic CDS (gray arrow) starts at the zero point. **D**. The pI value distribution of RefSeq and novel proteins. **E**. Bar chart of start codons used in RefSeq versus novel CDS, shown for four main categories and others. **F**. A novel CDS (66 aa), predicted as antitoxin MazE9, is shown in the IGV genome browser along with annotation sources (for simplicity, we show the most relevant ones) and peptide evidence (red). **G**. A subset of 6 orthologs of novels identified by Ribo-seq data in H37Rv and by proteogenomics here exhibit a signal of purifying selection in the clinical reference strains (right panel) that is comparable to that observed among RefSeq proteins (left panel).

The 37 orthogroups of novel proteins are shown in the context of additional predictions and evidence (**Fig. 5A, Supplementary Table S10**). They included 5 novels seen in all six strains (novel_1-5), 20 lineage-specific cases (15 for L1 including novel_9, _11, and _12 identified in all L1 strains, and 5 for L2, with novel_18 and _23 seen in all L2 strains) and 12 novels that were identified in a subset of strains from both lineages. A mirror plot of the highest scoring PSM is shown for each peptide of the novel proteins per strain (**Supplementary Files**). Most novels (86%) were conserved in higher taxonomic ranks (green boxes, **Fig. 5A**), and only 5 candidates were restricted to the MTBC. Similarly, for most novels (84%), the encoding genes were transcribed (**Supplementary Fig. S12**) and for 17/37 (46%), the respective gene was predicted to belong to an operon. Notably, our novels were enriched in basic and acidic proteins with more extreme pI values compared to RefSeq proteins: eight with a pI ≥ 11, six ≤ 5) (**Fig. 5D**). Furthermore, a meta-analysis of all start codons uncovered an enrichment of novel proteins starting with the alternative start codon CTG (**Fig. 5E**). A prediction of putative functions returned a Phyre2 hit with over 50% confidence for 12 novels (**Fig. 4C**), and with over 90% confidence for five of them. These included several interesting candidates: novel_3 (66 aa) is predicted as antitoxin MazE9 (**Fig. 5A**), and encoded upstream of a gene annotated as type II toxin-antitoxin system PemK/MazF family toxin, a genomic orientation common for type II toxin-antitoxin systems^58^. Novel_7 (271 aa) is predicted as pentapeptide repeat protein MfpA (>99% confidence), which was shown to contribute to resistance to fluoroquinolone (FQ) antibiotics in *Mycobacterium smegmatis*, where it protected DNA gyrase and decreased FQ-introduced DNA cleavages^59^. The novel CDS represents a second, longer protein present in addition to the annotated MfpA (183 aa) with limited sequence identity. Based on overlaps with annotated proteins (annotated function in brackets) on the same or opposite strand, additional interesting candidates could be novel_30 (siderophore RND transporter MmpL5: drug efflux), novel_19 (zinc metalloprotease HtpX: virulence factor), novel_17 (mycofactocin biosynthesis chaperone MftB: survival in acidic environment) and novel_14 (ImmA/IrrE family metallo-endopeptidase: stress response, associated with toxin-antitoxin systems). Interestingly, nine of the 37 novel orthogroups (24.3%) were predicted to encode an AMP, only one of which had a Phyre2 prediction (see **Discussion**). Finally, for 14/37 of our novel candidates (38%) a Ribo-seq signal had been detected for the corresponding H37Rv ortholog^42^, including three also detected by MS in that study, which have since been included in the RefSeq2024 genome annotation. We explored whether any of our novel CDS exhibited a similar signal of purifying selection as was observed for 90 of the 2,299 pervasively translated mRNAs in H37Rv, indicating they could represent functional CDS^42^. Evidence for such a signal, a pronounced positive bias in the G/C to A/T proportion between the third and second codon position (see **Methods**), is observed for all RefSeq proteins (**Fig. 4G**, left panel). Notably, a subset of six novels also had a significant G/C skew (based on Fisher’s exact test and same criteria as in^42^, (**Fig. 4G**, right panel), which was much less pronounced and not significant over all novel proteins (**Fig. 4G**, middle panel).

## Discussion

Our multiOMICS study on clinical reference strains of L1 and L2 illustrates that Mtb is an excellent system for genomic and proteogenomic investigations. Although short read-based assemblies dominate, a meta-analysis of complete Mtb genomes from NCBI RefSeq uncovered that only 7% are repeat-rich and difficult to assemble class III genomes (**Fig. 1A**). Overall, Mtb strains are thus -with the exception of the W/Beijing family of L2^53^ and selected L4 strains^60^ that harbor very long repeats-much simpler to assemble than those of virtually all other important bacterial pathogens^34^, including *Pseudomonas aeruginosa, Acinetobacter baumannii and typhoidal Salmonella,* with class III genome percentages of roughly 55, 69 and 98% (**Fig. 1C**).

Complete genomes are the optimal basis for comparative genomics, as they do not inflate pangenome estimates like short read assemblies^21^. Relying on long read data^7^, we de novo assembled complete genomes for six L1 and L2 strains, which has recently also been accomplished for larger sets of isolates^20^. The large core genome of the six reference strains (3,877 genes) and indication for convergent deletions (**Fig. 2C**) agrees with results of a much larger MTBC pangenome study of 339 curated genomes from L1-L9 and animal-adapted Mtb lineage strains^15^, which uncovered that evolution of MTBC is dominated by deletions. The availability of many complete genomes, ideally from clinical reference strains that cover the diversity within and between lineages, is critical to avoid false conclusions based on few reference genomes that miss regions variably present in isolates. Missed regions (genomic blind spots) are also relevant for individual genomes, and prevalent in short read assemblies. Due to their relevance for virulence, immune evasion, host pathogen interaction and as potential source for the development of new drugs and vaccines^12,13,17^, we report six complete catalogs of PE/PPE family members, including those missed in genomic blind spots, specific to either lineage, only present in the clinical reference strains and in the large repeat regions of L2 strains (**Table 1**, **Supplementary Table S6**). We release a Snakemake pipeline to comprehensively identify and accurately classify them for other complete Mtb genomes.

Using ddaPASEF as an example of a modern single injection proteomics method readily available in core facilities, we covered 66-68% of the theoretical proteome per strain and identified novel CDS from total cell extracts. We report all proteogenomic novelty as orthogroups in the categories novel CDS, longer and shorter CDS than annotated by Refseq and expressed pseudogenes, readily visualizing novels conserved in all six strains, plus lineage- and strain-specific cases (**Fig. 3**, **Fig. 5A, Supplementary Table S10**). To our knowledge, this is the first proteogenomics study i) on clinical Mtb reference strains and ii) that relies on the FDP estimate provided by entrapment^32^ to assess separately and stringently control the higher local FDR for novel protein identifications, a fundamental -and often not adequately accounted for-issue of proteogenomic studies^29,46,71^. In contrast to the global 1% FDR for the protein level searches from FDRBench for RefSeq, a much higher FDP-estimated local FDR among novel proteins was observed, i.e., between 11-23% (**Fig. 4A, D**). The Percolator score distributions of peptides from different subsets (RefSeq, decoy, novel hits below the threshold) were clearly separated from those of peptides implying novel identifications that either passed our ad hoc PSM and peptide filters^29,41^ (LC) or additionally the statistically robust entrapment based FDP estimation (HC). Reassuringly, the score distributions were very similar to that of RefSeq proteins (**Fig. 4B**). The ad hoc filters thereby ensured a substantially reduced FDR for novel proteins reported in our past proteogenomics studies. Combined, these observations support reporting all novels in the classes HC (novels that are also retained using an entrapment-based FDR threshold) and LC. Strikingly, all novels predicted by Prodigal were classified as HC and 80% of the Ribo-seq informed orthologs from H37Rv, underlining the value of creating smaller custom iPtgxDBs based on experimental data. Conversely, in silico ORFs had a significantly lower pass rate in the entrapment-based FDR filtering (43%, **Fig. 4C**). Novel hits with 2 peptides or more mostly belonged to HC identifications (91.8%), with that rate dropping to 33% for single peptide hits. Yet, as argued before^44,61^, they should be listed as potential candidates, ideally followed by independent validation, which was beyond the scope here. Together, these insights represent useful guidelines to create tailored iPtgxDBs with the public web server, e.g. overall smaller ones prioritizing high quality experimental datasets or more credible prediction sources, and for the downstream analysis of the search results.

Our results are consistent with reports that pseudogenization is an important driver of strain diversity in Mtb^57^ which lack HGT: among the 44 pseudogene orthogroups, 37 were grouped with RefSeq CDS of other strains and in 30 cases the annotated pseudogene was only found in one of the strains. Also, we did find *espJ* (Rv3878.1) annotated as pseudogene or missing in 2 L1 strains, respectively (N0072, N0153, **Fig. S7**), while *espK* (Rv3789c.1) was pseudogenized in strain N0155 (L2). Both genes are located in the RD1 region deleted in the attenuated live vaccine BGC^62^. Our results are also in line with recent recognition that strain-specific genetic and transcriptional diversity, beyond lineage, is relevant to understand disease phenotypes^1,63^.

SEPs were significantly enriched among all novel CDS (65%, **Table 2**, **Fig. 5B**). This is highly relevant given that missed short CDS can number in the hundreds for bacterial genomes. For example, 140 CDS below 50aa in *E.coli* were validated over roughly a decade^64^, several hundred identified in recent large Ribo-seq studies^65^, and also by the large number of recently identified families of conserved small proteins^25,66^. The substantial enrichment for a non-canonical start site (CTG) among our novel longer/shorter CDS candidates (**Fig. 5E**) may be relevant; at least in Enterobacteria, use of non-canonical starts had an unsuspected function in metabolic competition^67^. The prediction of functions for novels is difficult with only few having been characterized^66^. Nevertheless, we found an antitoxin in a genomic arrangement typical for type II toxin-antitoxins^58^ (**Fig. 5F**). Different basal levels of antibiotic resistance have been described^68^ and could possibly play a role also in the two lineages: we found a second, longer gene encoding a protein similar to MfpA (novel_7, 271aa), involved in drug resistance to FQ antibiotics^59^. An AlphaFold prediction of its structure closely resembles that of the shorter 183 aa form, suggestive of a similar function despite very low sequence similarity (**Supplementary Fig. S13**). Moreover, AMPs^69^ have been largely overlooked so far, but can play important roles in shaping microbiome composition^28^. Nine of the 37 novel orthogroups have a high AMP prediction score (**Supplementary Table S10**) and could represent interesting candidates for experimental follow up. Finally, the signal of purifying selection for 6 of our novels, as reported for a small subset of pervasively translated ORFs in H37Rv^42^, implies that they could be functional CDS.

Overall, the results of our improved, yet generic proteogenomics method illustrate that proteogenomics on clinical Mtb reference strains can uncover evidence for relevant novel genes even from unfractionated total cell extracts and with stringent FDR control. This study could be readily extended to covering all 10 human-adapted MTBC lineages. Future efforts should ideally also include membrane fractions to increase the number of novels observed, and/or integrate data from Ribo-seq approaches relying on start- and stop site enrichment. Combined with the complete PE/PPE gene catalogs reported here, proteogenomics has potential to identify new targets for the development of diagnostics, vaccines or therapeutics.

## Acknowledgement

This study was funded by the Swiss National Science Foundation (grant 197391 to CA, grants CRSII5_213514, 320030-227432 and 320030L-231163 to SG, and Ambizione grant PZ00P3_161435 to BC), Queen’s University Belfast startup funds to BC and grant 883582 from the European Research Council to SG. We thank Wolfgang Hess (University of Freiburg) for critical feedback on the manuscript.

## Competing Interests

The authors declare no competing interests.

## Notes

### Competing Interest Statement

The authors have declared no competing interest.

### Summary of Updates

We replaced the previous peptide-level entrapment-based false discovery rate estimations and filtering with the respective protein-level entrapment FDR estimations. This allowed us to better evaluate the FDR contol among novel proteins and to visualize this in the revised Figure 4 (plus 2 new Supplemental Figures). We also updated selected method sections.

