## Supplemental Information for "Stringent proteogenomic discovery of novel small proteins in *Mycobacterium tuberculosis* clinical reference strains"

### Supplement Contents

### Supplementary Methods

#### Genomic DNA extraction and de novo genome assembly

Genomic DNA (gDNA) for Illumina sequencing was extracted from cultures in the late exponential growth phase using the CTAB method^1^; gDNA for PacBio sequencing was prepared as described^2^. *De novo* genome assemblies of the six strains were created with long PacBio reads from an RSII device ((2-4 SMRT cells per strain; Polymerase binding P6 version (v.) 2, sequencing reagents v.4.0))^3^ and the assembly algorithm Flye (v.2.4)^4^, using length filtered subreads (> 1 kb, except for N0072, where greater 5kb was used) and an estimated genome size of 4.4 Mbp. The assemblies were polished by 2-4 iterations using Quiver from the SMRT Portal (v.2.3.0) and PacBio Reads (> 1 kb) until a single variant level was reached. To correct any remaining small assembly errors, data from paired end Illumina libraries (prepared using NEXTERA XT DNA Library Prep Kit (Illumina, San Diego, USA) and sequenced on an Illumina HiSeq2500 (Illumina, San Diego, USA) with 151 or 101 cycles at the Genomics Facility Basel) were mapped to the respective assembly using BWA MEM (v.0.7.17)^5^. FreeBayes (v.1.0.0; minimum alternate fraction: 0.5, minimum alternate count: 5) was run for 2-5 iterations until no further sequence corrections were observed. Manual start-alignment of the assemblies was set to 200 bp upstream of the *dnaA* gene. To verify the circularity and completeness of the *de novo* assembly, the filtered PacBio subreads were re-mapped to the circular chromosome using graphmap (v.0.5.2)^6^. Structural variations were called using Sniffles (v.1.0.7)^7^ and manually inspected in the Integrated Genome Viewer^8^. Quality parameters for assemblies were calculated by QualiMap (v.2.2.1)^9^. To detect potential short plasmids that could have been missed in the Flye assembly, Plasmid SPAdes (v.3.13.0)^10^ was run on the Illumina reads. Finally, assemblies were annotated with NCBI’s Prokaryotic Genome Annotation Pipeline (PGAP v.4.8)^11^. For selected genome characteristics, see **Supplementary Table S2**.

#### Comparison of complete genomes and Illumina assemblies

Illumina reads were trimmed with trimmomatic v0.39^12^ in paired-end mode (options illuminaclip: NexteraPE, leading: 3, trailing: 3, slidingwindow: 4:15 and minlen: 36), quality checked before and after trimming using FastQC v0.11.9 and assembled with SPAdes v3.15.2^13^ using default settings. The resulting contigs were mapped to the complete PacBio assemblies using minimap2 (version 2.17)^14^ with asm5 preset (0.1% sequence divergence); the output of the genomecov command from bedtools v2.30.0^15^, with bga and split options enabled) was then filtered to identify regions longer than 5 bases that were missing in the Illumina assemblies. Missing/partially covered genes were determined by finding overlaps between missing regions and annotated coding sequences (CDS) in the PGAP GFF file using bedtools’ intersect command (with -wo flag). CDS for which greater than 5 percent of the total length were absent from the Illumina assembly were considered to be partially missing. A blastn 2.10.1+ search with the CDS against all Illumina contigs removed a few false positives with split Illumina contig alignments. Hypergeometric tests were used to explore whether CDS annotated by PGAP as PE, PPE or transposase gene family members were enriched among missing CDS. Mapped Illumina contigs, gaps and missing genes were visualized together with sliding window means of the PacBio read coverage (reads longer than 5kb mapped with graphmap 0.5.2, window size 80 kb and step size 5 kb) using Circos v0.69.8^16^ (**Supplementary Fig. S2**). To show the increased coverage of the repeat region in the L2 strains, sliding window means of the PacBio read coverage were visualized in R.

#### Classification of PE and PPE gene family members

##### Prediction of protein families

For each strain, PE and PPE family genes were identified using three methods. The first unambiguous family assignment was selected in the following order: 1) The product description of the PGAP annotation was searched for a matching regular expression (regex), 2) a reciprocal best BLAST hit (RBBH) was used to match all annotated PGAP proteins per strain to a set of manually curated and annotated PE and PPE proteins from H37Rv^17^, and 3) the descriptions of PFAM domains identified by an InterProScan 5.59-91.0 analysis of the annotated PGAP proteins were searched with the same regex set. The combined approach identified 2-3 additional members on top of the PGAP annotation.	Proteins annotated by PGAP were assigned to the PE family if the product description contained the regex '(^|[^\w])PE($|[^A-Za-z])' and to the PPE family if it contained the regex '(^|[^\w])PPE($|[^A-Za-z])'. Rare cases where the product description contained both patterns lead to an ambiguous assignment to both families (below 2% of the set of PE/PPE genes). RBBHs between the manually curated list of H37Rv PE and PPE family proteins^17^ and all PGAP annotated proteins per strain were determined using BLASTp 2.12.0 (default settings, word size of 3). The best hit per protein was identified by the highest BitScore; if an RBBH could be identified, the family classification predicted by Ates was used for the matching protein. InterproScan 5.59-91.0 was used to analyze all annotated PGAP proteins per strain and protein families were assigned by parsing the description of domains identified by Pfam 35.0 analogously to the product description of the PGAP annotation. Predictions of transmembrane domains and signal peptides by TMHMM 2.0c and SignalP 4.1, respectively, were also extracted from the InterProScan results.

##### Phylogenetic prediction of sublineages

Proteins unambiguously identified as PE or PPE family proteins by the approach described above were subjected to a phylogenetic analysis in order to further classify them into one of five sublineages of the corresponding protein family as described^18^. For this, the n-terminal domains of the protein sequences were extracted and separately aligned per family and strain using ClustalW 2.1. For PPE proteins, 180 n-terminal amino acids (aa) were extracted, while for PPE proteins, 95 instead of 110 n-terminal aa were used, as this resulted in better resolved trees. Consensus trees based on 1000 bootstrapped neighbour-joining trees were calculated from the alignments using PAUP* 4.0a168^19^, collapsing branches with less than 50% support. Based on the consensus trees, the sublineages of proteins without a RBBH were predicted: Leafs without a RBBH in clades where all other existing RBBH were assigned to the same sublineage were assigned to this sublineage using the ete3 Python library v3.1.3, which was also used to visualize the trees (data not shown).

##### Prediction of motif-based subfamilies

A motif-based subfamily prediction was used for those cases where the protein had no RBBH from which the annotated subfamily for PE or PPE genes could be assigned. The four subfamilies were: PE-PGRS (a subfamily of PE genes with glycine-rich repeats with motifs GGA and GGN), PPE-PPW (characterized by the presence of P..P..W and GF.GT aa motifs in the first 10/30 aa in the C-terminus), PPE-SVP (proteins with a C-terminally located Serine-Valine-Proline sequence with the motif G..SVP..W) and PPE-MPTR (proteins with major polymorphic tandem repeats centered around an N.G.GN.G motif (N = Glutamine). The occurrence of these motifs was counted in each protein sequence and the protein was assigned to the corresponding protein family if the count exceeded a fixed threshold: For the PE-PGRS family at least 16 motif matches in the protein sequence were required, for PP-SVP 1 match, for PPE-MPTR 3 matches and for PPE-PPW 2 matches. The information from the H37Rv reciprocal best BLAST hit, the PE phylogenetic analysis and the pattern searches was incorporated into a final classification for each PE and PPE genes, which is provided as a separate Excel table (**Supplementary Table S6**) and summarized in **Table 1.**

#### Correlation of gene and protein expression

Illumina HiSeq 2500 reads for the six Mtb strains from^20^ (ENA Project PRJEB33274) were quality-checked using fastQC 0.11.9 and mapped to the respective complete genome using bwa-backtrack (v0.7.17) in single-end mode. The combined lists of annotated proteins and novel proteins detected by MS were converted to GFF format while only keeping the longest identified isoform of each annotation cluster; mapped reads per gene were counted using htseq-count (v1.99.2) with the stranded=reverse option enabled. Read counts were converted to transcripts per million (TPM), PSM counts divided by the protein length and the resulting values log2 transformed. Pearson’s correlation was calculated using all genes that were expressed at the transcript and protein level, and subsets highlighted (**Supplementary Fig. S12**).

### Supplementary Figures and Tables

#### Supplementary Table S1: Overview of six complete* genome assemblies of *M. tuberculosis* L1 and L2.

|  |  | ***De novo* genome assembly** | | | **(Partially) lacking in respective**  **Illumina assembly** | | | |
| --- | --- | --- | --- | --- | --- | --- | --- | --- |
|  | ***Mtb* strain** | **Genome size** | **Genes (pseudo-**  **genes)** | **CDS** | **No.**  **Scaffolds**  **/Contigs**** | **Base pairs**  **(%)** | **CDS (pseudo-genes)**  ******* | **CDS**  **≤ 100aa** |
| ***L1*** | ***N0072*** | 4,421,403 | 4,156 (134) | 4,105 | 368 | 135,793 (3.1) | 160(14) | 25 |
|  | ***N0153*** | 4,403,638 | 4,133 (143) | 4,082 | 593 | 33,419 (0.8) | 60 (6) | 12 |
|  | ***N0157*** | 4,409,145 | 4,148 (145) | 4,097 | 264 | 23,991 (0.5) | 44 (3) | 8 |
| ***L2*** | ***N0052*** | 4,409,093 | 4,158 (137) | 4,107 | 216 | 35,473 (0.8) | 57 (16) | 11 |
|  | ***N0145*** | 4,415,638 | 4,175 (130) | 4,124 | 218 | 26,809 (0.6) | 55 (8) | 12 |
|  | ***N0155*** | 4,415,309 | 4,170 (132) | 4,119 | 274 | 23,529 (0.5) | 43 (10) | 12 |

* Genomes were assembled using paired end Illumina reads^3^, allowing multiply mapping reads (found e.g. in gene families with many closely related members like PE and PPE family^17^). **The number of contigs/scaffolds of our six Illumina-based assemblies. ***CDS that lack 5% or more of the complete sequence are listed (pseudogenes are shown in brackets).

#### Supplementary Table S2. Selected genome features of six clinical *M. tuberculosis* reference strains.

|  | ***Lineage 1*** | | | ***Lineage 2*** | | |
| --- | --- | --- | --- | --- | --- | --- |
|  | ***N0072*** | ***N0153*** | ***N0157*** | ***N0052*** | ***N0145*** | **N0155** |
| Genbank acc. # | CP040689 | CP040690 | CP040692 | CP040688 | CP040693 | CP040691 |
| No. chromosom. (plasmids) | 1  (0) | 1  (0) | 1  (0) | 1  (0) | 1  (0) | 1  (0) |
| Genome size (bp) | 4,421,403 | 4,403,638 | 4,409,145 | 4,409,093 | 4,415,638 | 4,415,309 |
| G+C content (%) | 65.22 % | 64.67 % | 64.71 % | 65.15 % | 65.09 % | 65.06 % |
| Mean coverage PacBio | 154x | 405x | 354x | 380x | 366x | 347x |
| Mean coverage Illumina | 57x | 133x | 105x | 204x | 194x | 156x |
| Tandem repeat, approx. genome coordinates* |  |  |  | ~380 kb  Mbp  3.47 - 3.85 | ~240 kb  Mbp  3.47 - 3.71 | ~260 kb  Mbp  3.47 - 3.73 |
| No. of genes** | 4,290 | 4,276 | 4,293 | 4,295 | 4,305 | 4,302 |
| No. of CDS*** | 4,105 | 4,082 | 4,097 | 4,107 | 4,124 | 4,119 |
| No. of PE genes**** | 88 | 81 | 84 | 89 | 88 | 91 |
| No. of PPE genes**** | 62 | 65 | 59 | 59 | 61 | 60 |
| PE or PPE genes**** | 7 | 7 | 7 | 7 | 7 | 7 |
| No. of rRNA operons (5S, 16S, 23S) | 1, 1, 1 | 1, 1, 1 | 1, 1, 1 | 1, 1, 1 | 1, 1, 1 | 1, 1, 1 |
| No. of tRNA genes | 45 | 45 | 45 | 45 | 45 | 45 |
| No. of nc RNA genes | 3 | 3 | 3 | 3 | 3 | 3 |
| No. pseudogenes  (PE / PPE) | 134  (5 / 4) | 143  (17 / 1) | 145  (9 / 4) | 137  (6 / 9) | 130  (7 / 7) | 132  (4 / 8) |
| No. of frame-shifted CDS (fCDS) | 1 | 0 | 11 | 15 | 20 | 18 |

*region with higher read coverage; ** includes CDS, misc. RNA genes, pseudogenes and fCDS entries (last 2 rows); ***CDS = protein coding sequences including fCDS entries; **** PE, PPE or PE/PPE genes as annotated by RefSeq before our analyses, which uncovered additional members of these families (see results text, **Table 1** and **Supplementary Table S6** for more detail)

#### Supplementary Fig. S1. Meta-data for genome assemblies of bacterial pathogens from NCBI RefSeq.

**A**. Classification of 7,193 *M. tuberculosis* assemblies based on the sequencing technology/ies used: PacBio (dark pink), Oxford Nanopore Technologies (ONT) (dark teal), hybrid versions combining short and long read date types (see legend), Illumina and other short read technologies (light gray), and unknown (dark gray), i.e., where no metadata was specified by the submitters.

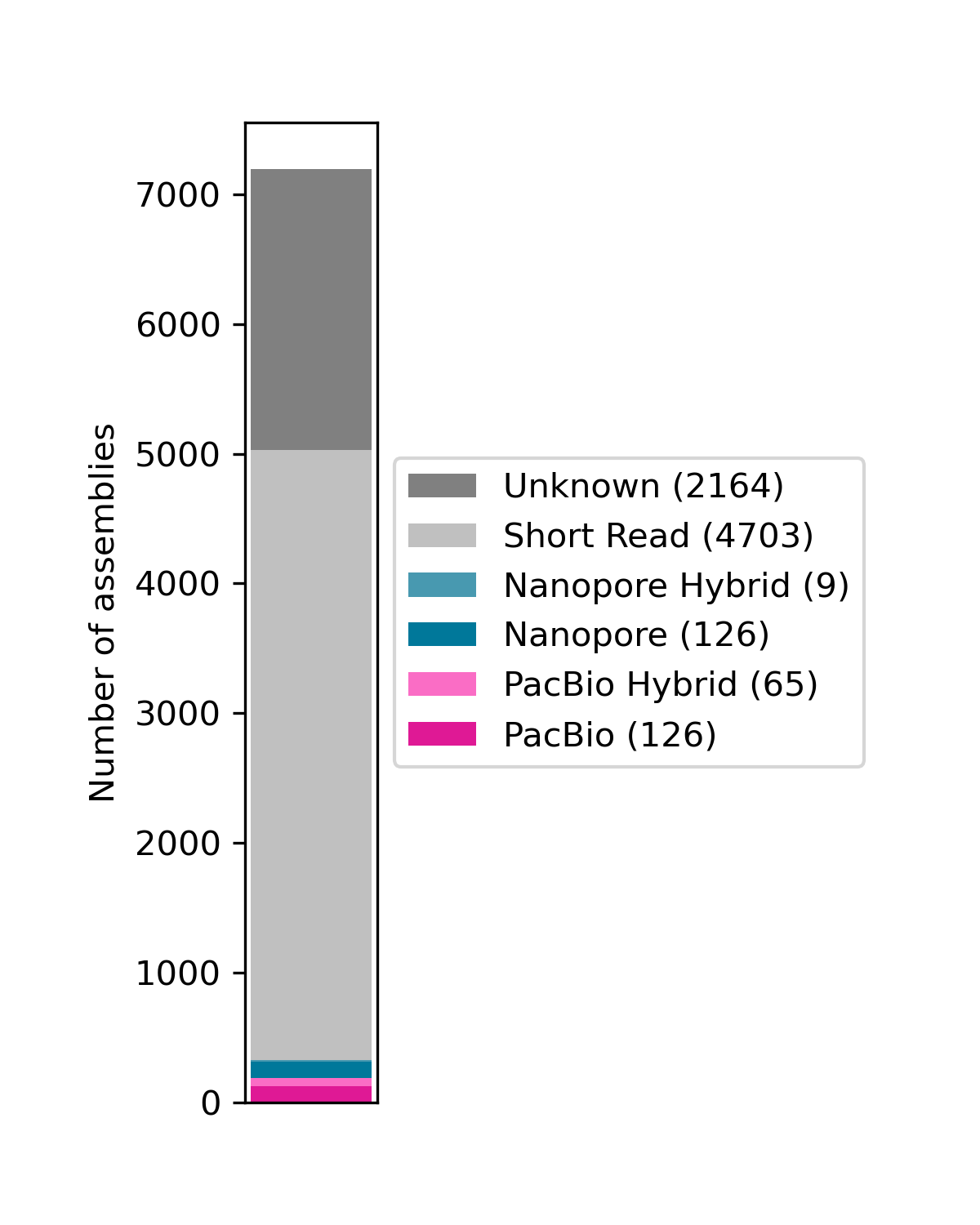

**B**. List of major bacterial pathogens and their genome assembly complexity. The list represents a combination of species from the WHO 2024 Bacterial Priority Pathogen List in the categories “critical” and “high priority” and from species, genera, or subsets of genera listed among 33 key bacterial pathogens that play a role in 11 infectious disease syndromes^21^. Below, we provide the genome assembly complexity classification of all publicly available complete genomes from NCBI RefSeq (as of Dec31, 2024) for these pathogens, the percentage of difficult to assemble class III genomes and an estimate of the number of deaths/year (except for Mtb, these numbers stem from the report mentioned^21^).

| **Species** | **Class 1** | **Class 2** | **Class 3** | **Total** | **ClassIII %** | **Deaths/Year** |
| --- | --- | --- | --- | --- | --- | --- |
| *Bacillus anthracis* | 120 | 0 | 7 | 127 | 5.5 | 2,000 |
| *Mycobacterium tuberculosis* | 399 | 0 | 30 | 429 | 7 | 1,250,000 |
| *Streptococcus pneumoniae* | 209 | 1 | 22 | 232 | 9.5 | 829,000 |
| *Neisseria gonorrhoeae* | 163 | 0 | 36 | 199 | 18.1 | 2,960 |
| *Enterococcus faecalis* | 222 | 0 | 51 | 273 | 18.7 | 220,000 |
| *Staphylococcus aureus* | 1,102 | 0 | 304 | 1,406 | 21.6 | 1,105,000 |
| *Yersinia pestis* | 0 | 57 | 18 | 75 | 24 | 1,500 |
| *Salmonella non-typhoidal* | 1,194 | 6 | 428 | 1,628 | 26.3 | 215,000 |
| *Campylobacter* spp | 504 | 0 | 186 | 690 | 27 | 123,000 |
| *Enterococcus* (other) | 81 | 5 | 34 | 120 | 28.3 | 100,000 |
| *Klebsiella* (other) | 269 | 20 | 128 | 417 | 30.7 | 53,900 |
| *Salmonella Paratyphi* | 17 | 0 | 8 | 25 | 32 | 23,300 |
| *Proteus* spp | 124 | 0 | 66 | 190 | 34.7 | 109,000 |
| *Serratia* spp | 174 | 3 | 96 | 273 | 35.2 | 100,000 |
| *Neisseria meningitidis* | 25 | 52 | 43 | 120 | 35.8 | 141,000 |
| *Morganella* spp | 49 | 0 | 29 | 78 | 37.2 | 5,510 |
| *Klebsiella pneumoniae* | 1,247 | 280 | 990 | 2,517 | 39.3 | 790,000 |
| *Enterococcus faecium* | 66 | 131 | 140 | 337 | 41.5 | 219,000 |
| *Citrobacter* spp | 208 | 26 | 169 | 403 | 41.9 | 54,100 |
| *Enterobacter* spp | 370 | 16 | 318 | 704 | 45.2 | 324,000 |
| *Shigella* spp | 11 | 95 | 100 | 206 | 48.5 | 113,000 |
| *Pseudomonas aeruginosa* | 408 | 4 | 509 | 921 | 55.3 | 559,000 |
| *Providencia* spp | 58 | 1 | 85 | 144 | 59 | 5,030 |
| *Escherichia coli* | 976 | 537 | 2,216 | 3,729 | 59.4 | 950,000 |
| *Listeria monocytogenes* | 162 | 0 | 260 | 422 | 61.6 | 14,900 |
| *Clostridioides difficile* | 72 | 0 | 117 | 189 | 61.9 | 33,200 |
| *Streptococcus other* | 368 | 13 | 824 | 1,205 | 68.4 | 518,000 |
| *Acinetobacter baumannii* | 223 | 12 | 533 | 768 | 69.4 | 452,000 |
| *Legionella* spp | 36 | 1 | 145 | 182 | 79.7 | 56,400 |
| *Vibrio cholerae* | 19 | 2 | 118 | 139 | 84.9 | 96,400 |
| *Aeromonas* spp | 28 | 14 | 264 | 306 | 86.3 | 21,300 |
| *Salmonella Typhi* | 2 | 0 | 128 | 130 | 98.5 | 182,000 |

#### Supplementary Fig. S2. Genome coverage of six *de novo* assembled clinical reference strains.

Coverage plot of PacBio reads (smoothed) for each of the six *de novo* assembled clinical reference strains from L1 (N0072, N0153, N0157) and L2 (N0052, N0145, N0155). A relatively uniform coverage (pink line) is observed for strains from L1 (upper panels). In contrast, the three coverage plots of L2 strains (blue line, lower panels) showed evidence for an unresolved, large repeat region of between 240-380 kb in length as indicated by the higher coverage. The start of the *dosR* operon is indicated by a black vertical line. The list of annotated genes contained in the respective repeat regions is available in **Supplementary Table S3**.

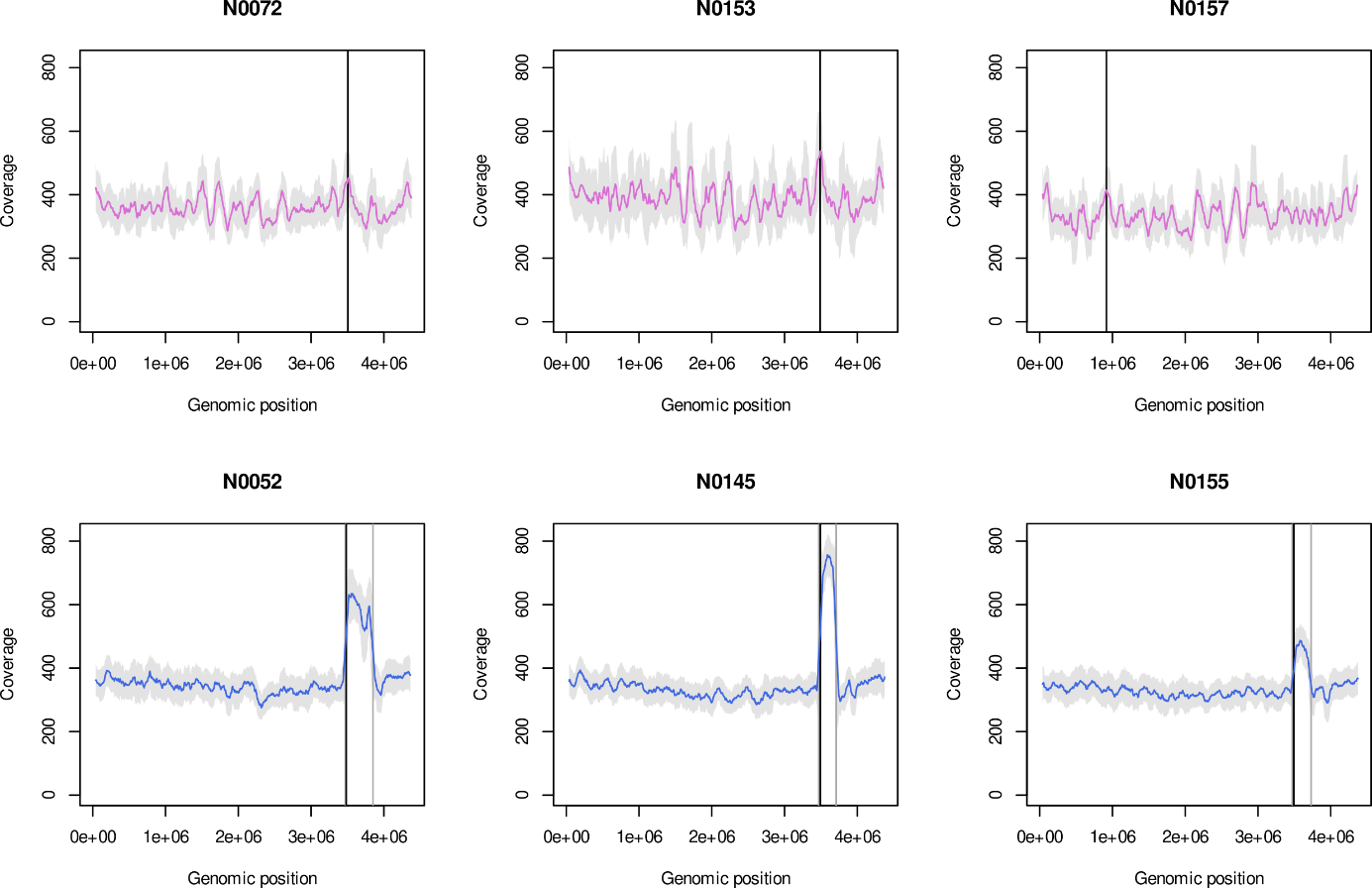

#### Supplementary Fig. S3. Genome assembly comparison.

For the four strains not shown in **Fig. 2B**, the outer rings show the complete genome (PacBio assembly), followed by the Illumina contigs (blue bars), the observed gaps (blue dashes), and a circle for genes missed (gray dashes) where three families were significantly enriched: PE genes (red dashes), PPE genes (green dashes) and transposons (purple dashes). The PacBio read coverage is shown in the innermost circle.

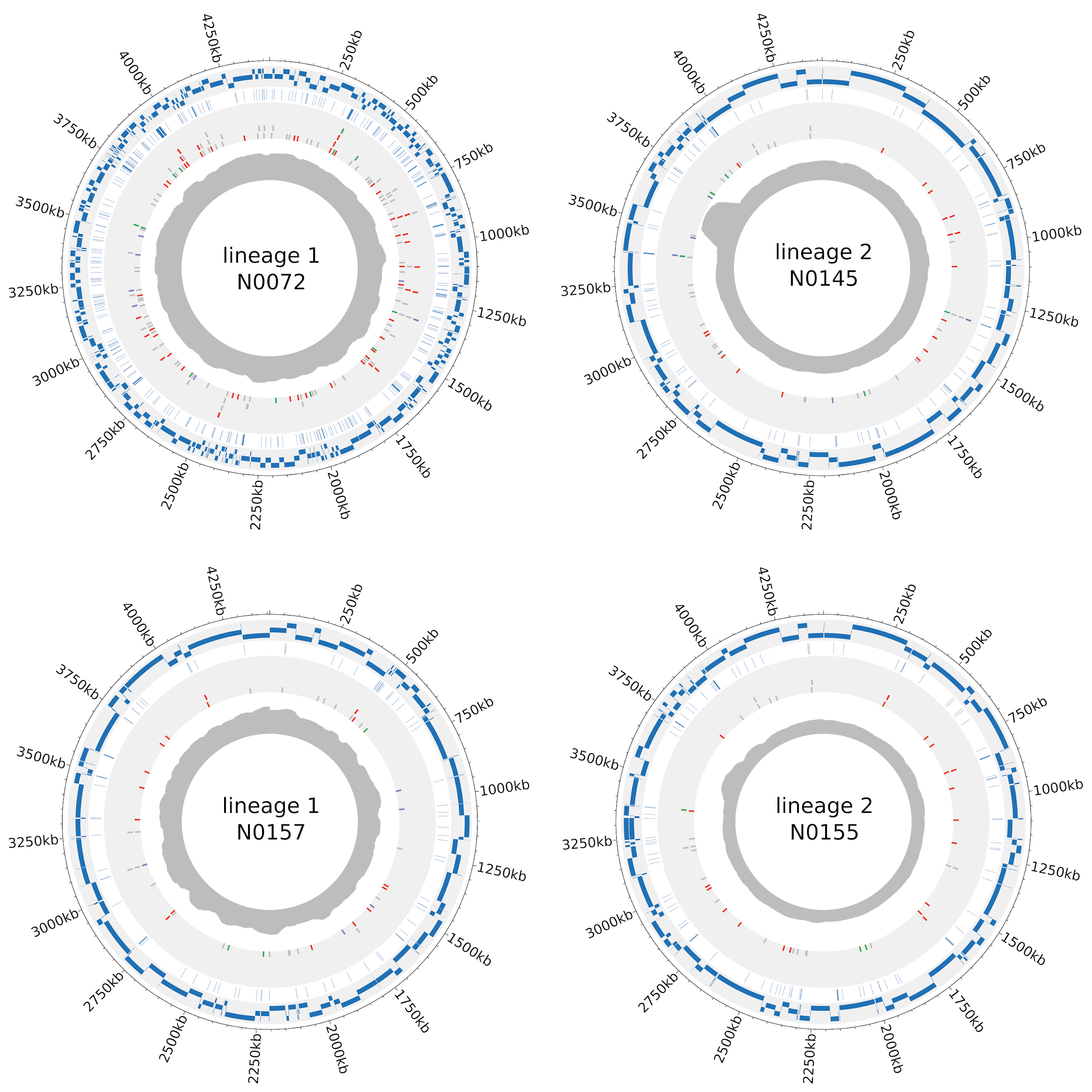

#### Supplementary Table S3. Genes in repeat regions with higher read coverage in L2 strains.

Overview of the genomic regions of *M. tuberculosis* L2 strains that exhibit a higher read coverage in the *de novo* genome assemblies and their annotated genes (NCBI’s PGAP), some of which include PE/PPE gene family members (**Table 1, Supplementary Table S6**). **See separate Excel file**.

#### Supplementary Table S4: Gene families enriched among CDS (partially) lacking in fragmented assemblies.

|  | **Complete genome** | | | **Missed in Illumina assembly** | | | **Hypergeometric P-values** | |
| --- | --- | --- | --- | --- | --- | --- | --- | --- |
|  | CDS | PE / PPE^#^ | Transp. | CDS | PE / PPE^#^ | Transp. | PE / PPE^#^ | Transp. |
| **N0072** | 4,105 | 159 | 44 | 160 | 63 | 7 | 1.02E-51 | 1.36E-03 |
| **N0153** | 4,082 | 155 | 42 | 60 | 27 | 5 | 9.51E-24 | 3.25E-04 |
| **N0157** | 4,097 | 152 | 54 | 44 | 19 | 5 | 1.05E-16 | 2.42E-04 |
| **N0052** | 4,107 | 157 | 61 | 57 | 24 | 4 | 3.04E-20 | 9.67E-03 |
| **N0145** | 4,124 | 158 | 64 | 55 | 23 | 6 | 2.55E-19 | 1.76E-04 |
| **N0155** | 4,119 | 160 | 62 | 43 | 23 | 0 | 2.68E-22 | 1.00E+00 |

^#^We grouped proteins annotated by PGAP as PE-family and PE-domain containing as well as PPE family and PPE-domain containing, excluding pseudogenes. For a more detailed classification into subgroups, see **Table 1** and **Supplementary Table S6**. Transp. = Transposons

#### Supplementary Fig. S4. Overlap of PE and PPE genes among 6 strains based on Panaroo.

The figure shows the results of the comparative genomic analysis using Panaroo, here specifically focusing on the PE (**A**) and PPE (**B**) gene families. Four PE genes are specific to L1, and 3 to L2 (**Supplementary** **Fig. S6A**). Among the PPE genes, we find 3 genes specific for L1, but no L2-specific cases. (**Supplementary** **Fig. S6B**). For more details, see **Supplementary** **Table S6.**

**A**

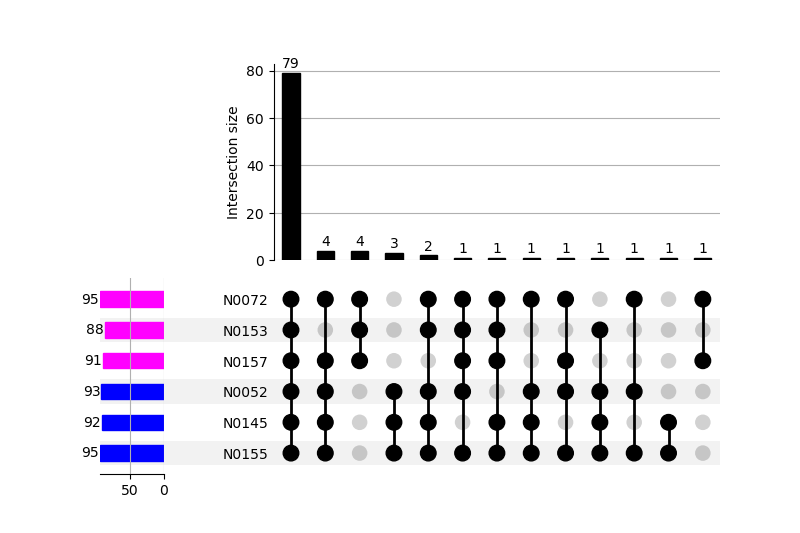

**B**

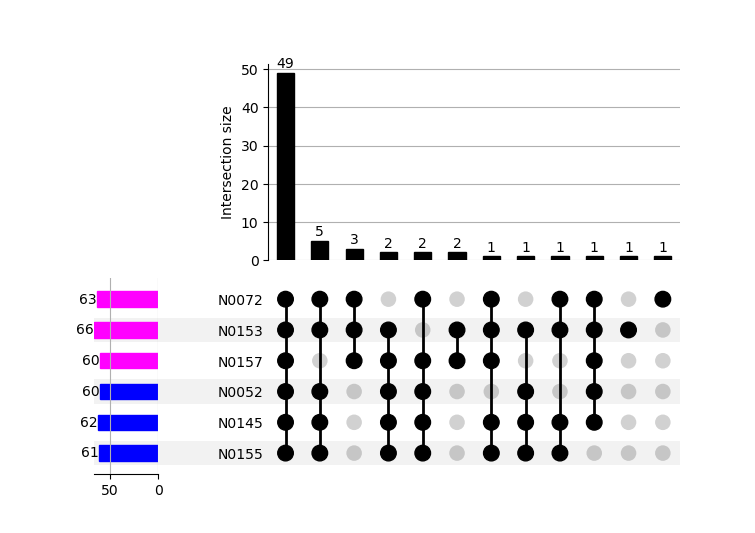

#### Supplementary Fig. S5. Configuration of the ppe38-71 locus in the six clinical reference strains.

The configuration of the *ppe38-71* gene locus was assessed and categorized based on a detailed analysis carried out previously^22^. All three L2 strains have an adjacent transposon gene (black arrow) that has been linked with a modern lineage; this gene is absent from L1 strains. The *ppe38-71* gene locus has been linked with the ability to secrete proteins of the PPE-MTPR and PE_PGRS families^22^. Proteins of these subfamilies are substrates of the type VII secretion system ESX-5, which have been shown to be relevant for virulence.

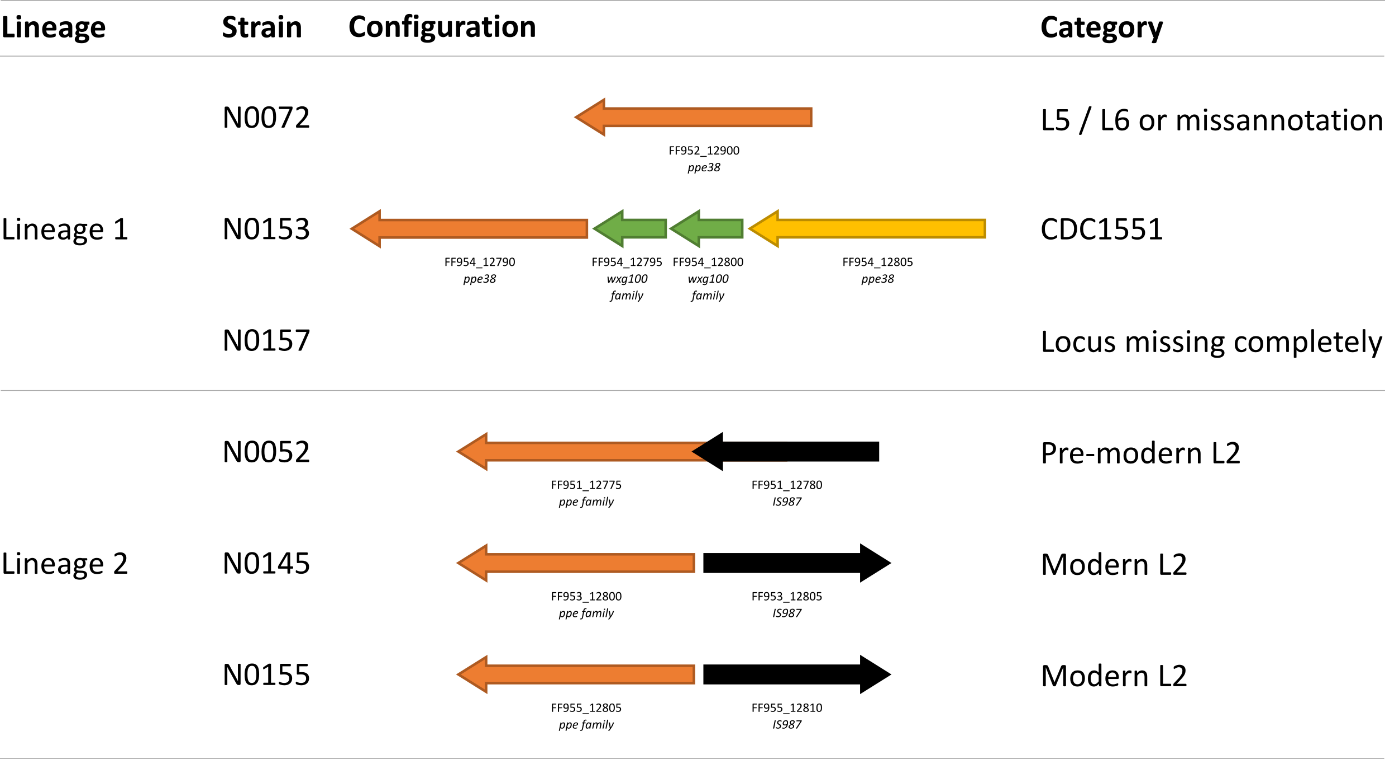

#### Supplementary Fig. S6. Percentage of unambiguous class 1a peptides for selected gene families.

**A**. A PeptideClassifier analysis of all RefSeq proteins of the respective strains indicated that the percentage of ambiguous peptides that do not allow to distinguish from which gene model an experimentally observed protein is expressed (class 3b peptides) is roughly 2-fold (except strain N0153) to 3.3 - 5.5 fold higher among proteins of the PE and PPE families, respectively, compared to that fraction for all RefSeq proteins. Overall, class 1a peptides dominate. **B**. The lower percentage of 1a peptides observed for both PE genes (t-test P-value < 0.0028) and PPE genes (t-test P-value < 9x E-9) is significant compared to the higher percentage of 1a peptides for all PGAP annotated proteins (Other).

**A**

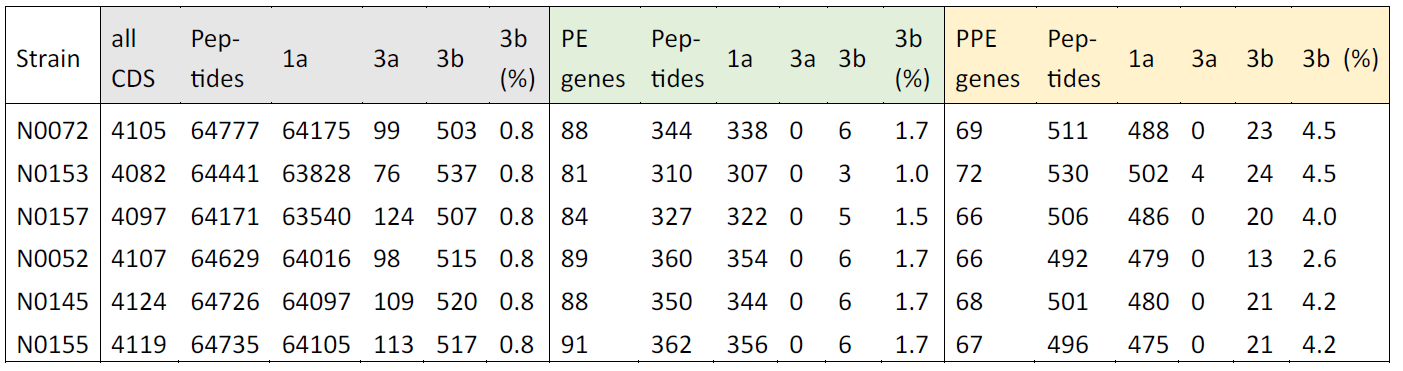

**B**

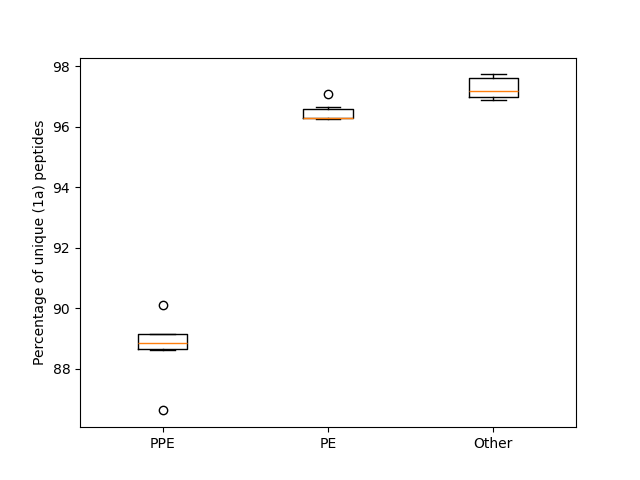

#### Supplementary Table S5. Result of the comparative genomics analysis of the six strains.

The conservation of the genes across the six strains is shown for each locus tag identifier and overall 4,198 gene clusters (see also **Fig. 2C**). Core genes: present in all six strains (3,877); strain-specific: annotated in only 1/6 strains (13); accessory: genes seen in a subset of two-five but not all strains (308). The accessory genes include L1-specific: genes conserved in lineage 1 strains (54); L2-specific: genes conserved in lineage 2 strains (33). **See separate Excel file**.

#### Supplementary Table S6. Classification of mycobacterial PE and PPE family genes into subcategories.

This table contains a worksheet for each of the six analyzed strains. Each worksheet has a list of PE and PPE family proteins identified by the PGAP annotation (columns G-L), a reciprocal best BLAST hit (RBBH) against a set of manually curated and annotated PE and PPE proteins from H37Rv^17^ (columns AA to AN), or a matching domain identified by InterProScan 5.59-91.0 (columns Q-U). Subfamilies were predicted based on the RBBH, and if no hit was found by a search of identifying motifs (columns V-Z). These family and subfamily predictions were consolidated into a final prediction (columns A-E) and sublineages were identified by a phylogenetic analysis of n-terminal domains, using the sublineage of the RBBH where present (column F). Since PE and PPE genes contain many repetitive sequences, the ambiguity of the tryptic peptides was evaluated using PeptideClassifier^23^ for each protein (columns N-O), listing peptides unique to each protein when they existed (column AO). Finally, proteins present in the repeat region of the lineage 2 strains (column M) and proteins that would have been missed in Illumina-only assemblies (columns K-L) were marked. For more detail, see the corresponding sections in the **Supplementary Methods**. **See separate Excel file**.

#### Supplementary Table S7. Proteins identified by a RefSeq search for six clinical reference strains.

| Strain | Identified | Coverage | Total SEP | Identified (SEP) | Coverage (SEP) | Prot. FDR |
| --- | --- | --- | --- | --- | --- | --- |
| N0072 | 2,749 | 67.0% | 536 | 186 | 34.7% | 0.52% |
| N0153 | 2,698 | 66.1% | 530 | 174 | 32.8% | 0.57% |
| N0157 | 2,699 | 65.9% | 532 | 167 | 31.4% | 0.45% |
| N0052 | 2,758 | 67.2% | 531 | 173 | 32.6% | 0.71% |
| N0145 | 2,789 | 67.6% | 533 | 178 | 33.4% | 0.58% |
| N0155 | 2,748 | 66.7% | 531 | 171 | 32.2% | 0.60% |

#### Supplementary Table S8. Proteomics evidence for genes missed in fragmented Illumina assemblies.

The overview below summarizes the search results against the respective RefSeq-based protein search DBs for the subset of proteins that were identified among the partially or entirely lacking protein coding genes (see Methods). These included between 6-10 proteins per strain, with the 57 proteins in N0072 representing an outlier that reflects the more fragmented assembly for this strain. Examples of relevant proteins are mentioned in the results; they include amongst others the ESAT-6 like protein EsxL, a SEP < 100 aa that was identified in 5 of 6 strains (N0072, N0153, N0157, N0145, N0155). **See separate Excel file**.

| Strain | Missing CDS | Missing CDS Id'ed* | %age |
| --- | --- | --- | --- |
| N0072 | 160 | 57 | 35.6 |
| N0153 | 60 | 10 | 16.7 |
| N0157 | 44 | 8 | 18.2 |
| N0052 | 57 | 6 | 10.5 |
| N0145 | 55 | 9 | 16.4 |
| N0155 | 43 | 6 | 14.0 |

#### Supplementary Fig. S7. Pseudogenes identified by proteogenomics and matching RefSeq CDS.

The overall 44 gene clusters that included expressed pseudogenes identified by our proteogenomics searches stood out: for many of the six clinical strains, a bona fide RefSeq annotated CDS existed (see legend), which matched an annotated pseudogene in a subset of the strains (34 of 44). This may indicate a process of pseudogenization^24^ in some Mtb clinical strains.

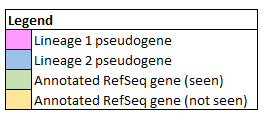

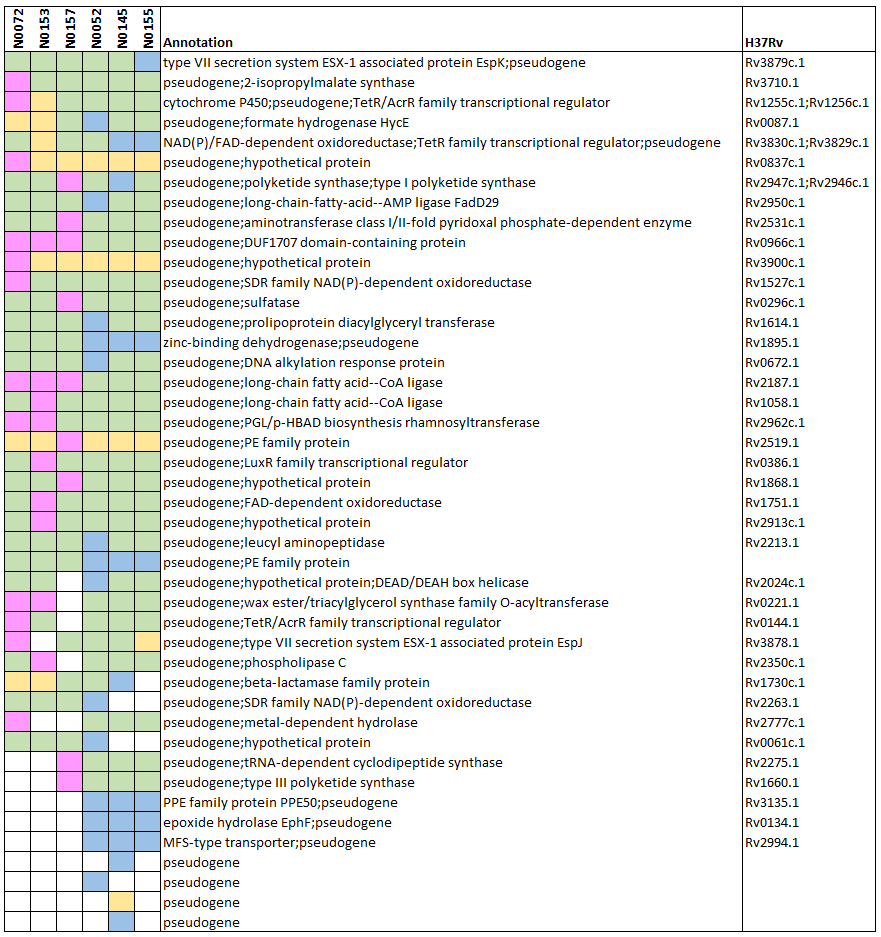

#### Supplementary Fig. S8. Identification of different protein start sites than annotated by RefSeq.

Overview of longer (**A**) and shorter CDS than annotated (**B**) identified by proteogenomics.

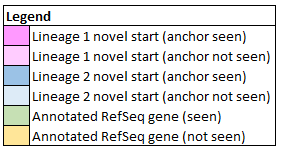

**A**

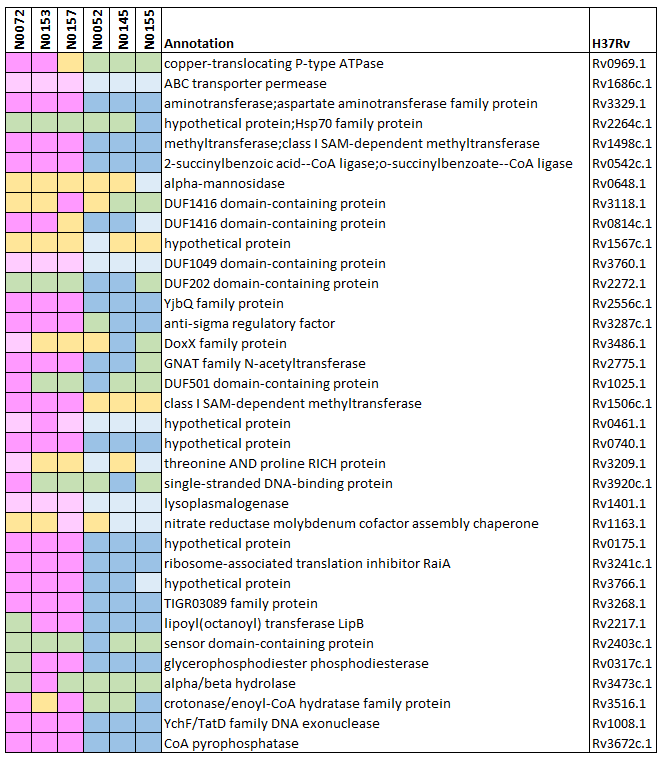

**A (continued)**

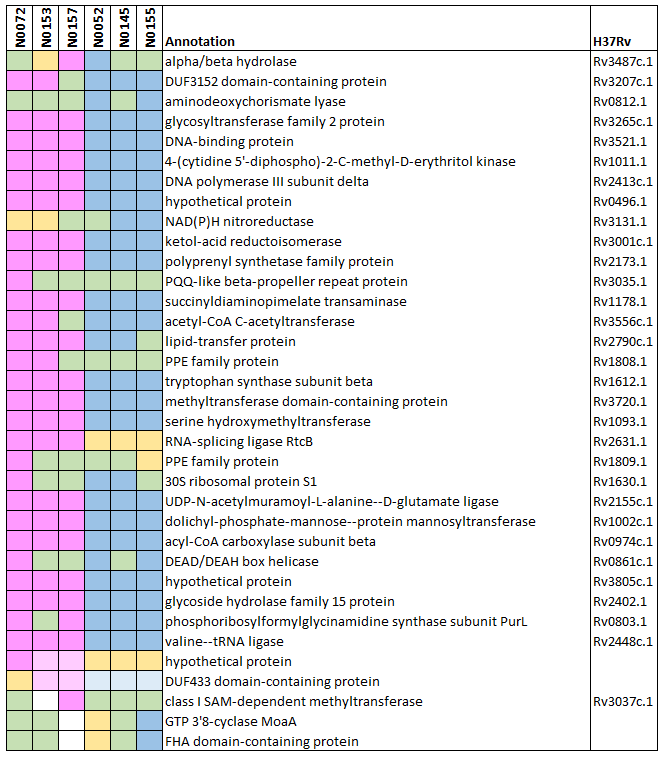

**B**

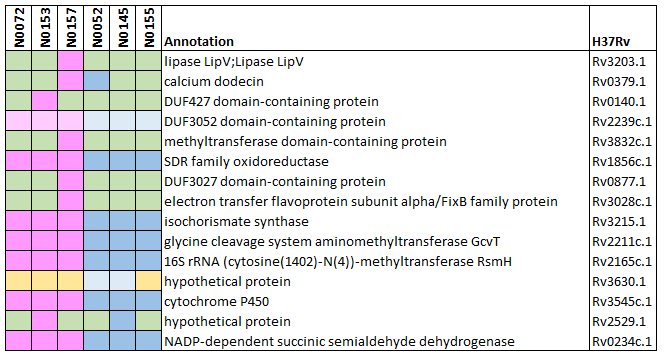

#### Supplementary Fig. S9. FDP curves of searches against standard iPtgxDBs.

Entrapment based false discovery proportion (FDP) curves for data before (left) and after applying the ad hoc filter (right) for i) all proteins (top row), ii) RefSeq proteins only (middle row), and iii) novel proteins (bottom row). Bands show the standard deviation between the six strains.

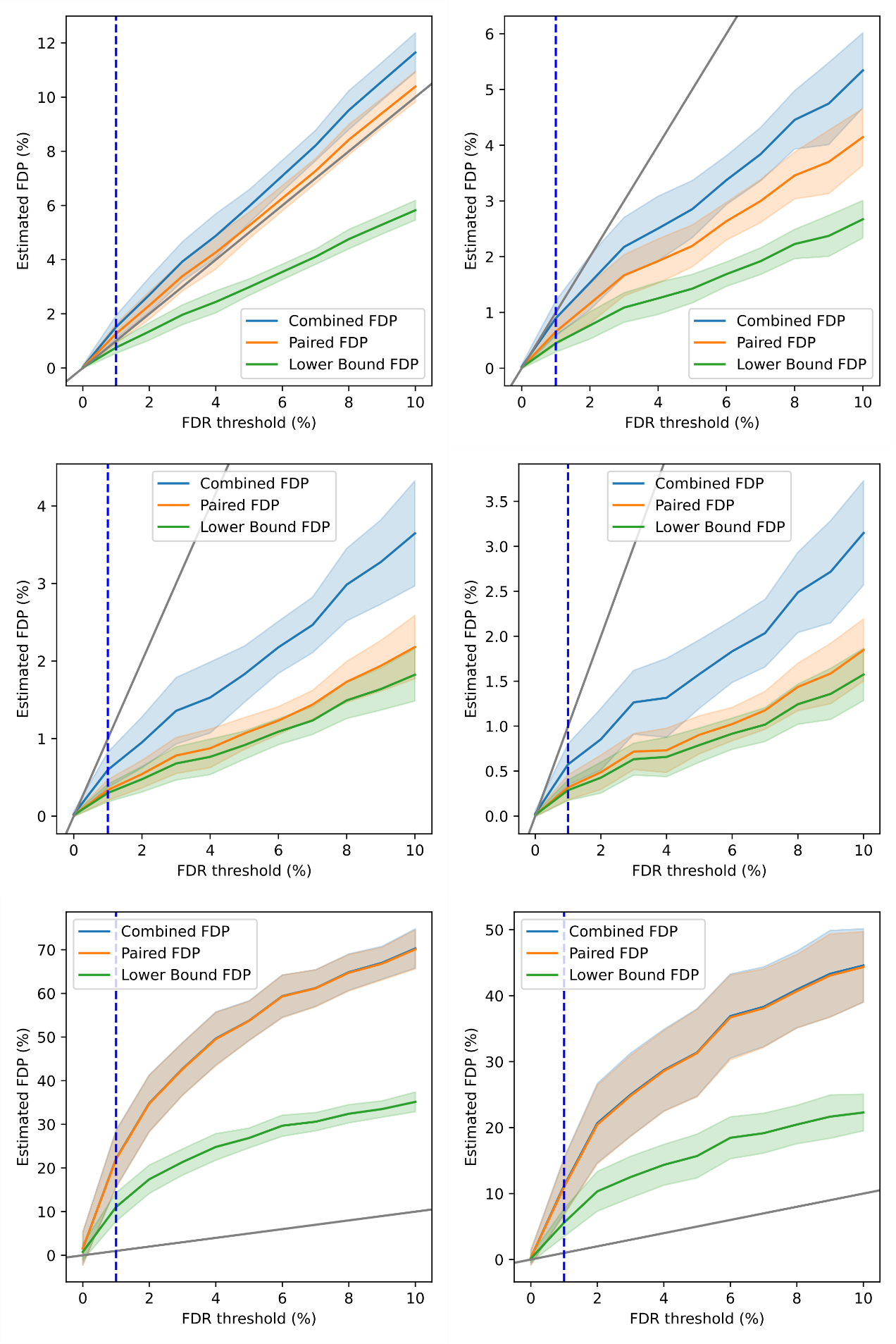

#### Supplementary Fig. S10. FDP curves of searches against custom iPtgxDBs.

Entrapment based FDP curves for data before (left) and after applying the ad hoc filter (right) for i) all proteins (top row), ii) RefSeq proteins only (middle row), and iii) novel proteins (bottom row). Bands show the standard deviation between the six strains.

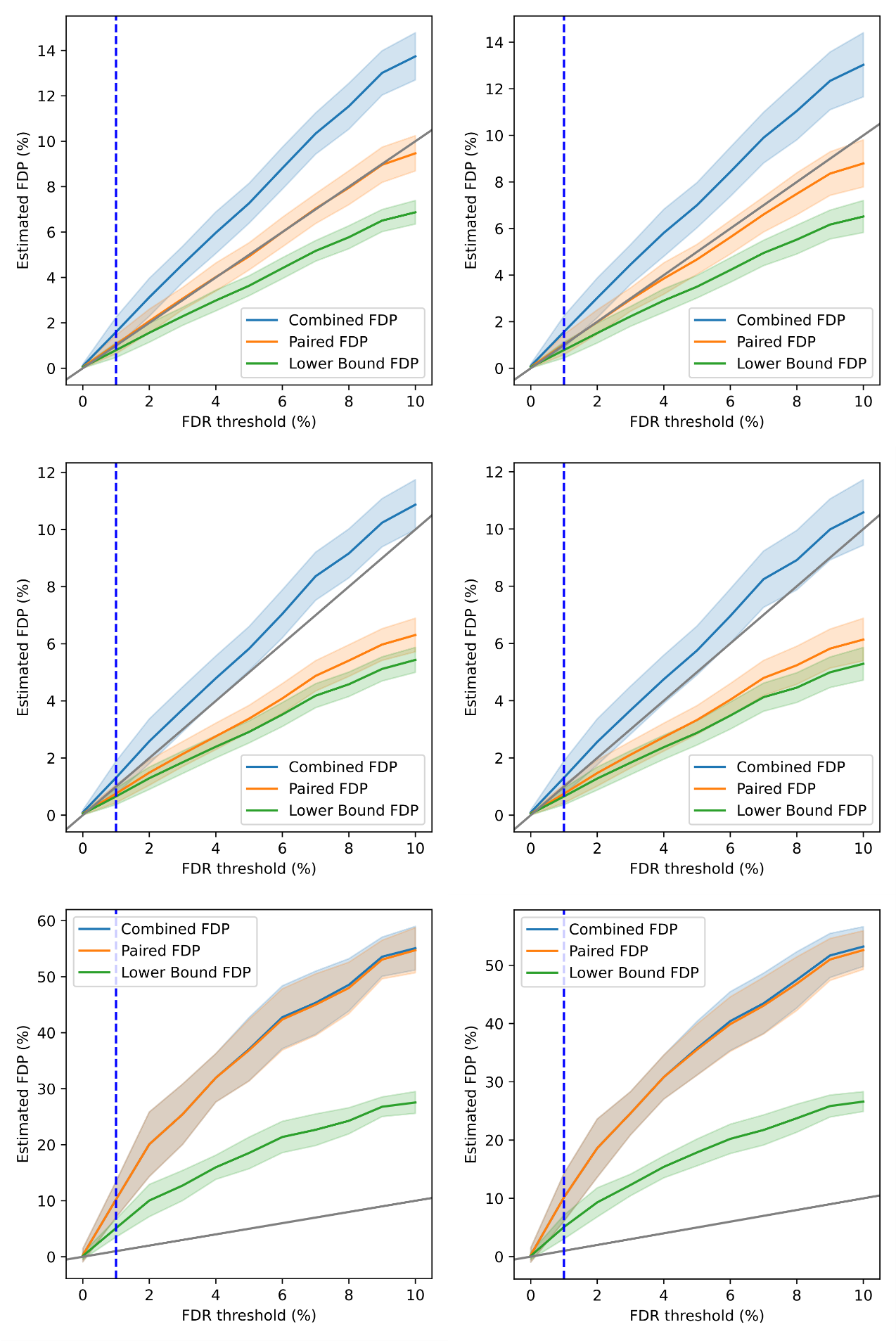

#### Supplementary Table S9. False discovery rate threshold selected based on entrapment filter.

Based on the estimated FDP for the subset of novels, new FDR thresholds were defined corresponding to an estimated FDP of 1%. These FDR thresholds shown below were then used to classify novel proteins into lower and higher confidence.

| **iPtgxDB** | **N0072** | **N0153** | **N0157** | **N0052** | **N0145** | **N0155** |
| --- | --- | --- | --- | --- | --- | --- |
| standard | 0.03% | 0.10% | 0.03% | 0.10% | 0.10% | 0.03% |
| custom | 0.20% | 0.20% | 0.24% | 0.07% | 0.30% | 0.10% |

#### Supplementary Fig. S11. Number of novel proteins identified per clinical reference strain.

The figure below shows the number of novel proteins detected in the searches against the standard and custom iPtgxDBs for each of the six strains and how many of these pass the two different quality filters that were applied: The ad-hoc filter is based on the number of PSMs and peptides (see methods for details) and has been used for all of our previous proteogenomics analyses while the entrapment-based FDP filter has been explored as an alternative method for the first time here. Proteins not passing any of the two filters (grey) or only the FDP filter (red) were discarded as putative false positives (also motivated by the results shown in **Fig. 4B**). Proteins passing the ad-hoc filter were categorized as lower quality identifications if they did not also pass the FDP filter (blue) and as higher quality identification if they passed both filters (purple). Overall, we thus report a stringently filtered subset as novel CDS.

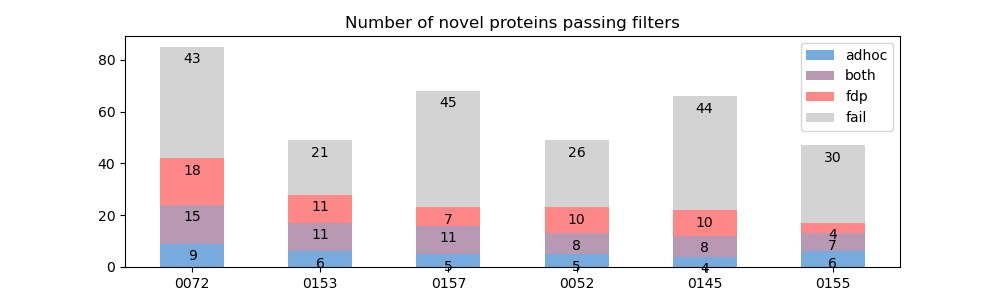

#### Supplementary Table S10. Pan-genomic consolidation of novel proteins including further bioinformatic analyses

The ad-hoc filtered novel proteins were consolidated into 37 pan-genomic orthogroups and further analyzed, including whether they were added to a newer RefSeq annotation or detected in two other studies (columns D-H), details on their proteogenomic detection (columns I-V) including entrapment filtering (columns W-AC) and whether they were also detected with transcriptomics (columns AD-AJ). Further, their conservation at different taxonomic levels (columns AK-AR), the overlap with RefSeq annotated proteins and the signal of selection based on codons not overlapping any annotated protein (columns AS-AV) was analyzed. For further insights into potential functions, a multitude of predictions were performed on the sequences of the novel proteins (columns AW-BM): structural similarity (Phyre2), function as an antimicrobial peptide (AMP Scanner), presence of known domains (eggNOG) and signal peptides (SignalP, LipoP), as well as transmembrane domains (TMHMM, Phobius), predicted subcellular localization (PSORTb) and inclusion in an operon (OperonMapper) were predicted. Finally, the physico-chemical parameters (columns BN-BQ) and the iPtgxDB identifiers for the identification in each strain (columns BR-BW) are provided. **See separate Excel file**.

#### Supplementary Fig. S12. Correlation of gene and protein expression data including novel proteins.

We analyzed RNA-seq data from^20^ as additional line of support for the 37 orthoclusters of novel proteins and 85 orthoclusters of longer or shorter CDS than annotated by RefSeq that we identified by our proteogenomics approach.

**A**. RNA-seq evidence indicates that the novel proteins are mostly well-expressed. The global correlation of gene (RNA-Seq) and protein expression (log2 values of (ddaPASEF observed PSMs divided by protein length)) data for all six clinical reference strains is plotted, as well as the respective Pearson correlation values (straight lines). RefSeq annotated genes/products are shown in light gray, extensions and reductions in blue and novel candidates including SEPs in red. Expressed pseudogenes were not considered for this plot.

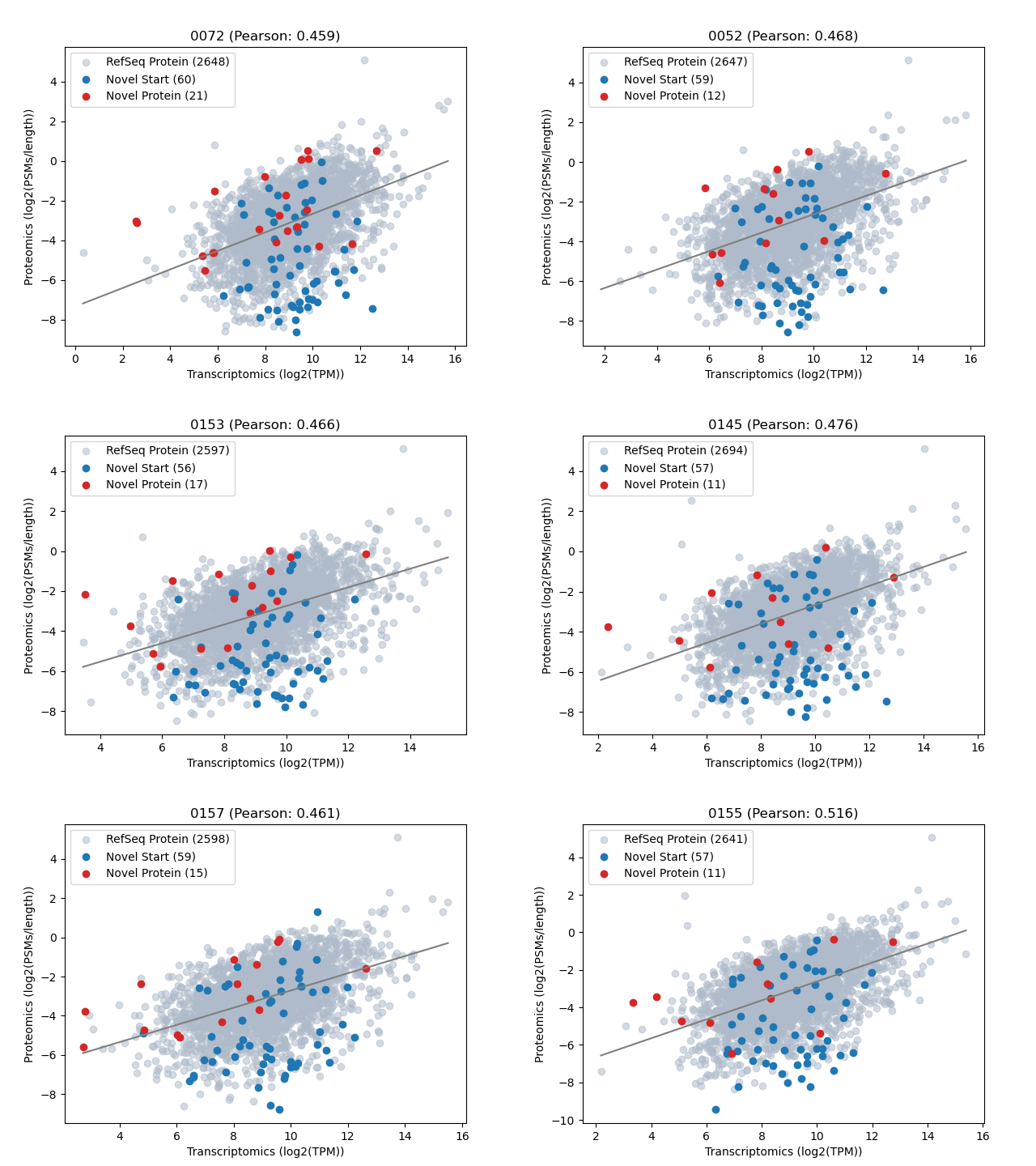

**B**. Density of log2 TPM expression data values with a focus on novel genes (blue crosses) and the subset of novel genes encoding small proteins below 100 aa (red crosses). Most of the genes encoding novel proteins are well expressed. Novel_6 (81aa) was identified in 5 strains and had the lowest transcript expression levels in 4/6 strains (N0072, N0153, N0157, N0155).

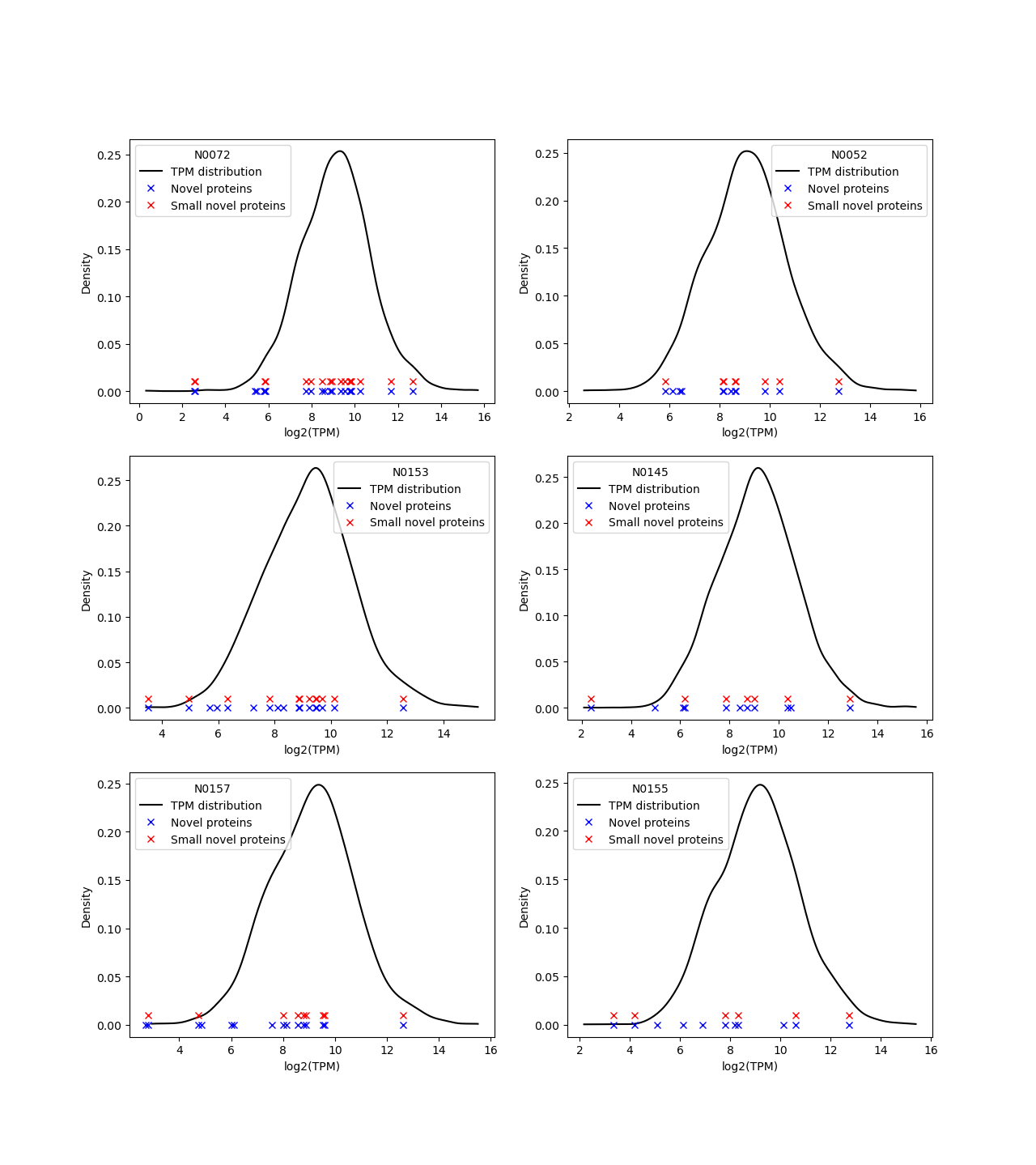

#### Supplementary Table S11. Summary of annotations integrated into the strain-specific iPtgxDBs.

We created iPtgxDBs as described previously^25^. We do not integrate pseudogenes in the annotation; instead, we handle them in a post-processing step. Entries were excluded if i) smaller than 6 aa, as these would not be detectable by MS (mostly extensions of RefSeq annotations), ii) entries whose internal start site (reductions) would not be distinguishable from a shorter proteoform of the longer PGAP annotation (if they both would start with a Methionine), and iii) if they were annotated as pseudogenes by PGAP and no other annotation source predicted a gene with the same stop.

**L1 strains:**

**N0072**

iPtgxDB composition summary

| Annotation source | Clusters | Entries | Added clusters | Added entries | Extensions | Reductions |
| --- | --- | --- | --- | --- | --- | --- |
| RefSeq (refseq) | 4,239 | 4,239 | 4,105 | 4,105 | 0 | 0 |
| Prodigal (prod) | 4,055 | 4,055 | 237 | 674 | 307 | 185 |
| ChemGenome (chemg) | 4,780 | 4,780 | 1,900 | 3,411 | 1,454 | 60 |
| *in-silico* ORFs (orf) | 69,900 | 137,352 | 63,658 | 113,328 | 49,469 | 224 |
| TOTAL | - | - | 69,900 | 121,518 | 51,230 | 469 |

Entries excluded from iPtgxDB

| Reason | RefSeq | Prodigal | ChemGenome | in-silico ORFs | TOTAL |
| --- | --- | --- | --- | --- | --- |
| too small | 0 | 97 | 73 | 2,393 | 2,563 |
| all-pseudo | 134 | 0 | 0 | 0 | 134 |
| indistinguishable start | 0 | 329 | 14 | 28 | 371 |

**N0153**

iPtgxDB composition summary

| Annotation source | Clusters | Entries | Added clusters | Added entries | Extensions | Reductions |
| --- | --- | --- | --- | --- | --- | --- |
| RefSeq (refseq) | 4,225 | 4,225 | 4,082 | 4,082 | 0 | 0 |
| Prodigal (prod) | 4,044 | 4,044 | 241 | 700 | 310 | 208 |
| ChemGenome (chemg) | 4,772 | 4,772 | 1,895 | 3,400 | 1,450 | 58 |
| *in-silico* ORFs (orf) | 69,581 | 136,753 | 63,363 | 112,762 | 49,200 | 226 |
| TOTAL | - | - | 69,581 | 120,944 | 50,960 | 492 |

Entries excluded from iPtgxDB

| Reason | RefSeq | Prodigal | ChemGenome | in-silico ORFs | TOTAL |
| --- | --- | --- | --- | --- | --- |
| too small | 0 | 95 | 70 | 2,384 | 2,549 |
| all-pseudo | 143 | 0 | 0 | 0 | 143 |
| indistinguishable start | 0 | 318 | 14 | 27 | 359 |

**N0157**

iPtgxDB composition summary

| Annotation source | Clusters | Entries | Added clusters | Added entries | Extensions | Reductions |
| --- | --- | --- | --- | --- | --- | --- |
| RefSeq (refseq) | 4,242 | 4,242 | 4,097 | 4,097 | 0 | 0 |
| Prodigal (prod) | 4,073 | 4,073 | 263 | 700 | 302 | 198 |
| ChemGenome (chemg) | 4,731 | 4,731 | 1,885 | 3,412 | 1,464 | 65 |
| *in-silico* ORFs (orf) | 69,816 | 137,346 | 63,571 | 113,201 | 49,390 | 263 |
| TOTAL | - | - | 69,816 | 121,410 | 51,156 | 526 |

Entries excluded from iPtgxDB

| Reason | RefSeq | Prodigal | ChemGenome | in-silico ORFs | TOTAL |
| --- | --- | --- | --- | --- | --- |
| too small | 0 | 108 | 69 | 2,394 | 2,571 |
| all-pseudo | 145 | 0 | 0 | 0 | 145 |
| indistinguishable start | 0 | 316 | 13 | 26 | 355 |

**L2 strains:**

**N0052**

iPtgxDB composition summary

| Annotation source | Clusters | Entries | Added clusters | Added entries | Extensions | Reductions |
| --- | --- | --- | --- | --- | --- | --- |
| RefSeq (refseq) | 4,244 | 4,244 | 4,107 | 4,107 | 0 | 0 |
| Prodigal (prod) | 4,072 | 4,072 | 251 | 684 | 297 | 190 |
| ChemGenome (chemg) | 4,762 | 4,762 | 1,879 | 3,397 | 1,468 | 53 |
| *in-silico* ORFs (orf) | 69,825 | 137,240 | 63,588 | 113,078 | 49,302 | 211 |
| TOTAL | - | - | 69,825 | 121,266 | 51,067 | 454 |

Entries excluded from iPtgxDB

| Reason | RefSeq | Prodigal | ChemGenome | in-silico ORFs | TOTAL |
| --- | --- | --- | --- | --- | --- |
| too small | 0 | 111 | 67 | 2,332 | 2,510 |
| all-pseudo | 137 | 0 | 0 | 0 | 137 |
| indistinguishable start | 0 | 328 | 15 | 23 | 366 |

**N0145**

iPtgxDB composition summary

| Annotation source | Clusters | Entries | Added clusters | Added entries | Extensions | Reductions |
| --- | --- | --- | --- | --- | --- | --- |
| RefSeq (refseq) | 4,254 | 4,254 | 4,124 | 4,124 | 0 | 0 |
| Prodigal (prod) | 4,079 | 4,079 | 241 | 681 | 305 | 186 |
| ChemGenome (chemg) | 4,763 | 4,763 | 1,876 | 3,392 | 1,464 | 53 |
| *in-silico* ORFs (orf) | 69,970 | 137,481 | 63,729 | 113,367 | 49,466 | 197 |
| TOTAL | - | - | 69,970 | 121,564 | 51,235 | 436 |

Entries excluded from iPtgxDB

| Reason | RefSeq | Prodigal | ChemGenome | in-silico ORFs | TOTAL |
| --- | --- | --- | --- | --- | --- |
| too small | 0 | 113 | 65 | 2,336 | 2,514 |
| all-pseudo | 130 | 0 | 0 | 0 | 130 |
| indistinguishable start | 0 | 331 | 15 | 18 | 364 |

**N0155**

iPtgxDB composition summary

| Annotation source | Clusters | Entries | Added clusters | Added entries | Extensions | Reductions |
| --- | --- | --- | --- | --- | --- | --- |
| RefSeq (refseq) | 4,251 | 4,251 | 4,119 | 4,119 | 0 | 0 |
| Prodigal (prod) | 4,082 | 4,082 | 250 | 687 | 302 | 188 |
| ChemGenome (chemg) | 4,762 | 4,762 | 1,874 | 3,392 | 1,465 | 56 |
| *in-silico* ORFs (orf) | 69,945 | 137,515 | 63,702 | 113,367 | 49,462 | 224 |
| TOTAL | - | - | 69,945 | 121,565 | 51,229 | 468 |

Entries excluded from iPtgxDB

| Reason | RefSeq | Prodigal | ChemGenome | in-silico ORFs | TOTAL |
| --- | --- | --- | --- | --- | --- |
| too small | 0 | 111 | 65 | 2,355 | 2,531 |
| all-pseudo | 132 | 0 | 0 | 0 | 132 |
| indistinguishable start | 0 | 328 | 14 | 22 | 364 |

#### Supplementary Fig. S13. Predicted structure of H37Rv MfpA and the longer novel CDS identified (strain N0072).

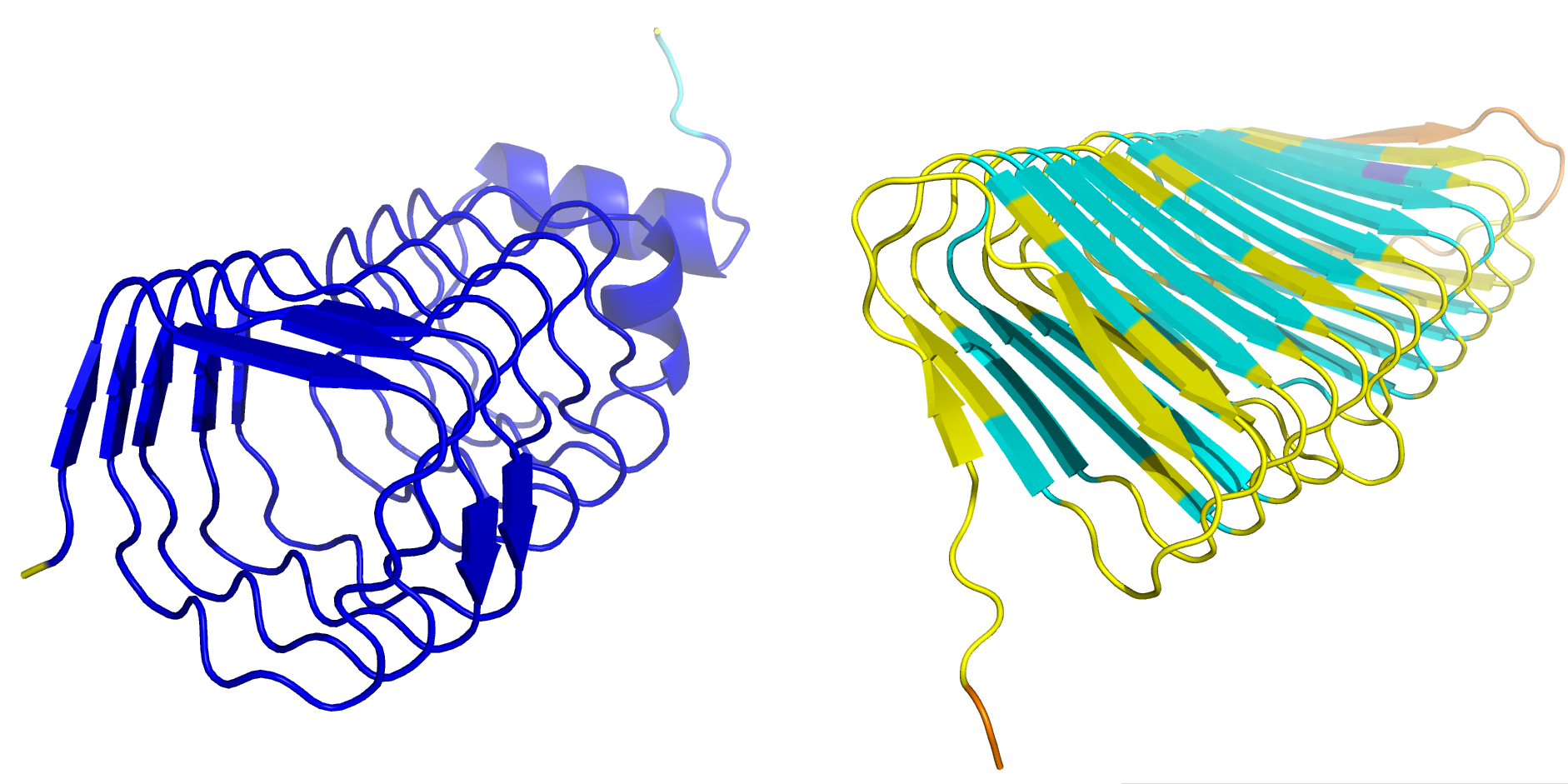

Predicted 3D protein structures of MfpA from H37Rv downloaded from UniProt (Rv3361c, left) and of the longer novel CDS identified in 5 of the 6 clinical reference strains calculated with AlphaFold server version 2025.07.05 (novel_7, here from strain N0072, right panel; **Fig. 5A**). The structures resemble each other quite closely, despite a very low amino acid sequence similarity.

### Supplementary Files

For each of the six strains, a PDF file is provided containing mirror plots of the best spectrum for each peptide identifying a novel protein. The best experimentally detected spectrum was selected according to Percolator score and hyperscore and is shown in the top half of the plot. Where possible, the matching *in silico* predicted spectrum for the same peptide (calculated with the Prosit_2023_intensity_timsTOF^26^ model using the Koina server^27^) is shown in the bottom half. In cases where no prediction was possible, the reason is stated instead. Fragment ion intensities shown on the y-axis were normalized and their m/z values are shown on the x-axis. Details shown in the header consist of the peptide sequence including modifications and the name of the novel protein. The second row shows the precursor charge, collision energy used, the m/z value of the precursor and the difference between theoretical and actual precursor mass in Dalton. The third row shows the Percolator score and hyperscore as calculated by Philosopher, followed by the spectral angle and Pearson’s correlation coefficient between observed and theoretical spectrum as calculated in^26^. Modifications in the peptide sequence are indicated by M[Ox] for oxidation of methionine and [Ac]- for N-terminal acetylation.
